# Globins in the marine Annelid *Platynereis dumerilii* shed new light on hemoglobin evolution in Bilaterians

**DOI:** 10.1101/789214

**Authors:** Solène Song, Viktor Starunov, Xavier Bailly, Christine Ruta, Pierre Kerner, Annemiek J.M. Cornelissen, Guillaume Balavoine

## Abstract

**Background:** How vascular systems and their respiratory pigments evolved is still debated. While many animals present a vascular system, hemoglobin exists as a blood pigment only in a few groups (Vertebrates, Annelids, a few Arthropod and Mollusk species). Hemoglobins are formed of globin sub-units, belonging to multigene families, in various multimeric assemblages. It was so far unclear whether hemoglobin families from different Bilaterian groups had a common origin.

**Results:** To unravel globin evolution in Bilaterians, we studied the marine Annelid *Platynereis dumerilii,* a species with a slow evolving genome. *Platynereis* exhibits a closed vascular system filled with extracellular hemoglobin. *Platynereis* genome and transcriptomes reveal a family of 19 globins, nine of which are predicted to be extracellular. Extracellular globins are produced by specialized cells lining the vessels of the segmental appendages of the worm, serving as gills, and thus likely participate in the assembly of the giant hexagonal bilayer hemoglobin of the worm. Extracellular globin mRNAs are absent in smaller juvenile, accumulate considerably in growing and more active worms and peak in swarming adults, as the need for O_2_ culminates. Next, we conducted a Metazoan-wide phylogenetic analysis of globins using data from complete genomes. We establish that five globin genes (stem globins) were present in the last common ancestor of Bilaterians. Based on these results, we propose a new nomenclature of globins, with five clades. All five ancestral stem-globin clades are retained in some Spiralians, while some clades disappeared early in Deuterostome and Ecdysozoan evolution. All known Bilaterian blood globin families are grouped in a single clade (clade I) together with intracellular globins of Bilaterians devoid of red blood.

**Conclusions:** We uncover a complex “pre-blood” evolution of globins, with an early gene radiation in ancestral Bilaterians. Circulating hemoglobins in various bilaterian groups evolved convergently, presumably in correlation with animal size and activity. However, all hemoglobins derive from a clade I globin, or cytoglobin, probably involved in intracellular O_2_ transit and regulation (clade I). The Annelid *Platynereis* is remarkable in having a large family of extracellular blood globins, while retaining all clades of ancestral Bilaterian globins.

## Background

The exchanges of gas, nutrient and waste relying on diffusion are impaired when body size and tissue thickness increase. Most animals develop at least one type of circulatory system to circumvent this limitation. Vascular systems are diverse, representing different solutions to the same purpose in animals with varied body plans.

One major function of the blood vascular system is performing gas exchanges, bringing dioxygen to the tissues and taking back waste products (eg. CO_2_). To perform this respiratory function efficiently, many species use respiratory pigments that bind dissolved gases in a cooperative way and considerably increase their solubility in the blood or hemolymph. A general picture of the nature of circulating respiratory pigments used in Bilaterians (Figure 1) suggests a complex evolutionary history.

**Figure 1:**
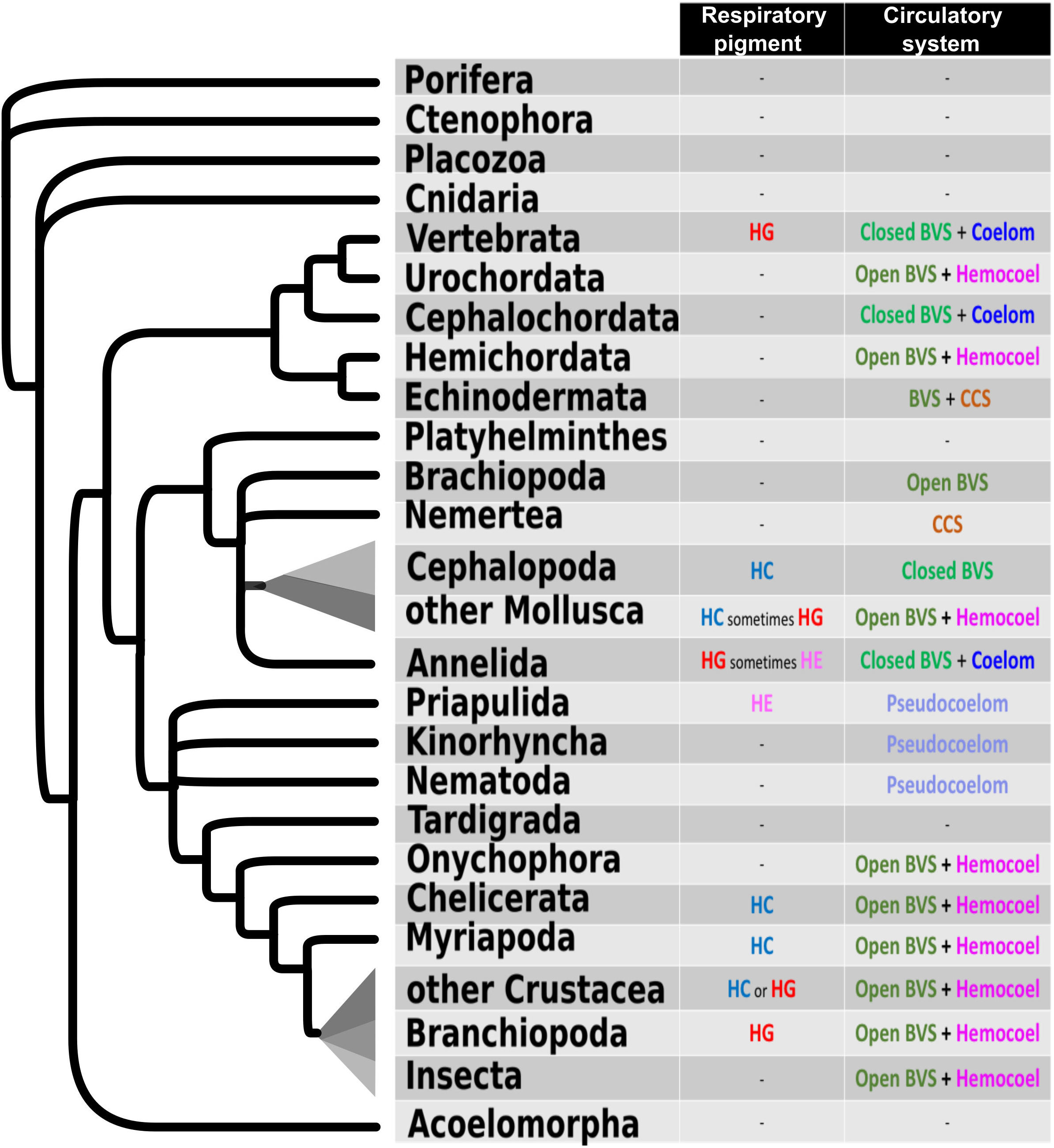
Distribution of circulatory systems and respiratory pigments in the Metazoan consensus tree,. Hb: hemoglobin. Hc: hemocyanin. Hm: hemerythrin. BVS: blood vascular system. CCS: closed coelomic system. Some species have no known respiratory pigments despite having a circulatory system or at least a fluid filled cavity (Nematodes, Echinoderms, Urochordates, Cephalochordates). It is assumed that in these species, either gases diffuse freely in thin layers of tissues or freely dissolved in the hemolymph. Some groups have circulating hemoglobins (“red blood”). The status of these hemoglobins is very diverse. They can be extracellular hemoglobins dissolved in the blood, as in many Annelids. They can be contained in red cells as in the Vertebrates or in some Annelids such as Capitellids (11). Some groups have circulating dissolved hemerythrins (“pink blood”), like Priapulids, Brachiopods and some Annelids (Sipunculidae, Magelonidae) (Mangum et Kondon 1975; Klippenstein 2015; Zhang et Kurtz 1991). Mollusks and many arthropods have circulating dissolved hemocyanins (“blue blood”). Circulating respiratory pigments of different types are generally not present together in the blood, suggesting that the recruitment of each type of pigment for the respiratory function occurred multiple times independently in the evolution of Bilaterians (74).

Among the circulating respiratory pigments, hemoglobins in particular display a complex situation and the evolution of globins is still not understood in depth. Globins are ancient proteins present in all groups of living organisms (1). They are characterized by a unique molecular structure, the “globin fold” made of eight alpha helices which shelters a heme prosthetic group, itself responsible for binding diatomic gases, such as O_2_ or NO. The globin motif domain is sometimes found in composite proteins that have evolved by fusion of pre-existing protein domains but all respiratory “hemoglobins” derive from proteins with a stand-alone globin motif. The oxygen transport function is well described in specific groups (Vertebrates, Annelids) but the globin superfamily is prevalent across the animal kingdom and the functions of globins are diverse and still poorly known (2). Many phylogenetic studies have focused on the hemoglobins of Mammals and Vertebrates in general, starting with the seminal work of Zuckerkandl and Pauling (1962). A smaller number of studies have focused on the evolution of Metazoan globins (4–7). These studies have shown that Vertebrate hemoglobins and the related myoglobins are only a small branch in a vast tree of animal globins, some of which are used for respiratory functions but many others have still unknown functions.

Some studies have highlighted the ancientness of neuroglobins, the first globins discovered in the Vertebrate neurons (8). These globins have homologues in groups as distantly related as Cnidarians and Sponges. Crucially, expressions of neuroglobins in Mammals, Cnidarians and the Acoel *Symsaggitifera* (7) all occur in neural or sensory cells. Neuroglobins are proposed to be the most ancient clade of globins occurring in animals. In a number of Bilaterians, hemoglobins are suggested to be derived from these ancestral neuroglobins by independent, convergent functional divergence (7).

Another study (9) has identified globin X, a molecule present in Teleost fishes and Amphibians but not in Amniotes, as an ancient globin clade. Globin X are expressed in neural cells as well in these Vertebrates (10). Another Panmetazoan clade of globins but missing in all Vertebrates, called X-like, is also identified (4). The same study showed that many of these Globins X or X-like carry a prototypical N-terminal acylation site (myristoylation and palmitoylation) suggesting that these proteins are linked to membranes in the cell. The authors proposed that Vertebrate hemoglobins could derive from molecules that were initially membrane bound.

These phylogenetic studies however have incomplete samplings of species and molecules. Gene sequences are identified from animals whose complete genome was not available at the time of the study and therefore represent only a subset of the diversity that must exist in Metazoans. A larger sampling of complete Metazoan genome sequences is now available and allows an update.

In this work, we explored the globin content of the genome of the marine Annelid *Platynereis dumerilii*. Many Annelids, including *Platynereis*, are remarkable for the similarities of their blood vascular system (BVS) with the Vertebrate BVS. Annelid BVS are usually closed. Annelid blood vessels, devoid of endothelium, are located in between spacious pairs of coelomic cavities in each segment and have a metameric organisation along the trunk of the animal. The blood of many Annelids is red, containing a high concentration of respiratory hemoglobin. In contrast to Vertebrates, this circulating hemoglobin is often extracellular instead of enclosed in red blood cells although in rare cases, some Annelids have circulating red blood cells with intracellular hemoglobin (11). The extracellular globin structure is very different from the hemoglobin of Vertebrates. Instead of a heterotetramer of globin sub-units, many Annelids possess a giant hemoglobin molecule, originally referred to as “erythrocruorin”, organised in a hexagonal bilayer of no less than 144 individual globin peptides (12–14), assembled with the help of linker proteins. *Platynereis* possesses red blood indicating that it is composed of dissolved hemoglobin. Despite its status of emergent model animal for evolution and development studies (15–17), the composition of the blood of *Platynereis* has not been yet characterized, although it can be inferred that they possess the Annelid extracellular giant hemoglobin. We identified the multigenic family encoding for the globins in *Platynereis* publicly available sequenced genomes and transcriptomes. In particular, we found a family of 9 extracellular globin genes, of which are found in a chromosomal cluster. These genes presumably code for the subunits that take part in the making of *Platynereis* giant extracellular respiratory pigment. We combined *in situ* hybridization and electron microscopy 3D reconstructions to identify and characterize the cells that produce these extracellular globins. We show that the morphology of these cells is indeed consistent with hemoglobin producing activity. We describe the changes of locations and intensities of this activity along the life cycle of the animal. The globin-producing cells are first found in lateral vessels appearing in between the segments. Then, in growing worms, they appear around the vessels inside parapodia, the gill-like appendages on which the worm relies for respiration. The extracellular globin gene expression intensifies as juveniles grow, culminates at the onset of sexual maturation and then collapses at the end of maturation process, as the worm is approaching sexual reproduction and the end of its life.

We analyse the evolutionary relationships of *Platynereis* globins at the metazoan-wide phylogenetic scale, using a sampling of Metazoans for which completely sequenced genomes are available. We wanted to put to the test the idea that animal hemoglobins are derived convergently from "neuroglobin-like" proteins, involved in a neuronal function (7). We show that a diversified set of at least five globins, the “stem globins”, of unknown functions, existed in the Bilaterian ancestor. We show that all families of Bilaterian blood globins (i.e. with O_2_ transport function) and their associated gene clusters arose convergently by repeated duplication events and concomitant functional recruitment from a single specific, cytoplasmic stem-globin, known as "cytoglobin" in Vertebrates (18) and not from a neuroglobin. This cytoglobin also exists, single or in a small gene family, in most Bilaterians devoid of red blood.

## Results

### Identification of globin genes in the Platynereis genome and cluster mapping

We identified in the transcriptomes and genome of *Platynereis* 19 different genes coding for proteins with a single globin motif (Figure 2A). Nine of these genes code for a peptide with a secretion signal at their N-terminus, (*Pdu Egb-A1a*, *-A1b*, *-A1c*, *-A1d-α*, *-A1d-β*, *-A1d-γ*. *-A2*, *-B1*, *-B2*), making them likely to represent extracellular globins (Additional file 1). Six of these genes, distinct from the predicted extracellular globins, code for peptides that are likely to be membrane-bound, thanks to N-terminal acylation sites. All 19 sequences were identified both in the genome assemblies and in the transcriptomes, indicating that they are not cross species contaminations. In addition, all sequences were mapped to different loci in the latest assembly of the *Platynereis* genome (Figure 2B and Additional file 2). They therefore represent genuine paralogues instead of divergent alleles. Six of the nine potential extracellular globins (*Pdu Egb-A1d-α*, *-A1d-β*, *-A1d-γ*, *-A2*, *-B1*, *-B2*) are comprised in a chromosomal cluster (Figure 2B) and two other closely related genes (*Pdu Egb-A1a*, *-A1c*), form a tandem pair in a distinct contig. All 10 other genes, coding for putative intracellular globins, as well as the extracellular *Pdu Egb-A1b* are found as isolated genes on rather large contigs.

**Figure 2:**
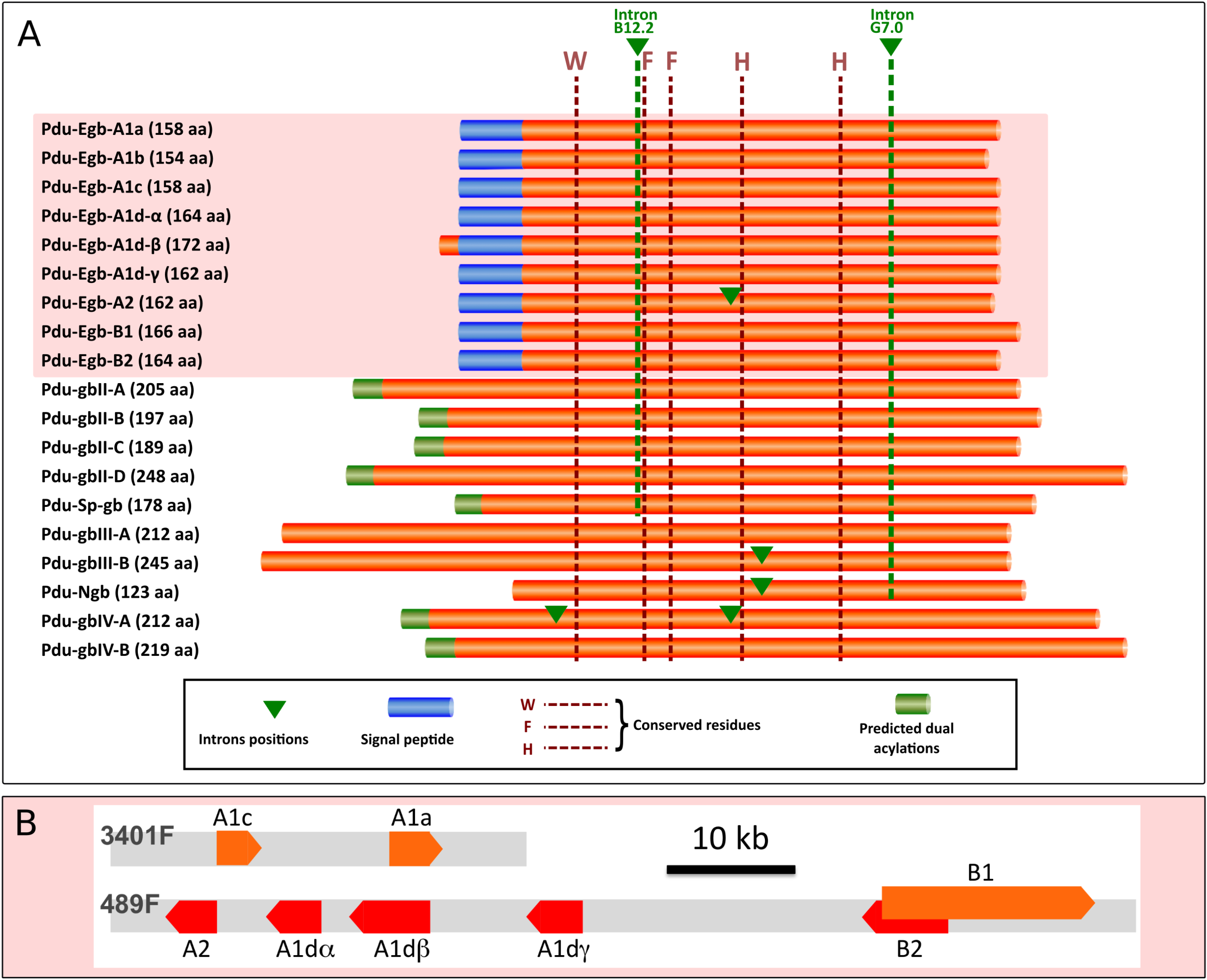
The globin genes of *Platynereis dumerilii*. A - Schematic representation of aligned primary structures of globin genes identified in *Platynereis* transcriptomes and genome. Five invariant amino acid positions provide solid anchor sites for precise homology alignments. Small Alignment gaps are not represented. The predominantly conserved introns positions B12.2 and G7.0 are marked by green stapled lines, Additional introns positions are shown with green arrowheads. Predicted N-terminal signal peptides and acylations are represented as shown. B - Schematic representation of the two globin gene clusters found on *Platynereis* scaffolds. Orange and red block arrows indicate the length and strand position of the genes. Each gene is interrupted by 3 - 4 introns. Gene *Egb-B1* partially overlaps with gene *Egb-B2*, by a small exon located inside one of the other gene introns.

### Phylogenetic analysis of the Platynereis globins at the Metazoan scale

To understand better how the origins of the multiple globin genes present in *Platynereis* genome, we carried out a phylogenetic analysis of Metazoan globins. Except for extracellular globins of some Annelid species, we used exclusively species for which a complete genome sequence is available. We chose species representative of all major Metazoan clades (phyla). We designed a screening technique with a concatenated globin probe and a low E-value cut-off in order to detect even the most divergent globins. We decided however not to treat in this study the globin domains that are part of multidomain proteins, such as the androglobin of Vertebrates. We also did not take into account the globins with multiple serially arranged globin folds as occurring for example in Echinoderms (19,20). We included the Crustacean *Daphnia pulex*. In *Daphnia*, 11 proteins have a double globin domain organization. As it suggested multiple tandem duplications of an ancestral didomain globin, we chose to separate the N- and C-terminal globin domains in each of these peptides and to treat them as individual OTUs in our trees. Our complete genome surveys uncovered a highly variable number of globin sequences in the different Metazoan species, ranging from none in the Ctenophore *Pleurobrachia bachei* and the Rotifer *Adineta vaga*, to 32 in the Annelid *Capitella teleta* (additional file 3). We also searched all Metazoan genomes for the specific linker proteins that are associated with the assembly of the giant hexagonal bilayer of Annelid erythrocruorin. These proteins are only present in the four Annelid species that have also extracellular globins (additional file 3).

We conducted a maximum-likelihood analysis of all 272 proteic sequences (sequence alignment in additional file 4; complete tree in additional file 5; simplified version in Figure 3). A major issue in phylogenetic analyses is to determine the root of the tree, as this directly impacts all the evolutionary interpretations that can be deduced from the tree. In this particular analysis, there are no obvious molecules that could serve as outgroups for globins inside the Metazoans. We also decided not to use globins from other Eukaryotic groups, as we were concerned that using very divergent outgroups with a small number of alignable positions could result in an artifactual rooting of the tree. Instead of determining where the root of the tree could possibly be, we decided to deduce where it could not be. To this goal, we took into account the solid groupings found in the unrooted tree, as indicated by their aLRT scores, and their composition in species to determine which groupings are natural clades, which we describe in details below. We also excluded a root position within well supported mono-species clades. The root of the tree that is represented in figure 3 (and additional file 5) is thus chosen arbitrarily among the positions that we consider possible based on that criterium. We looked for groupings that would indicate pan-Metazoan or pan-Bilaterian genes, because they contain globins that are derived from a large sampling of Metazoans species and do not contain sub-trees that are themselves composed of a large panel of Metazoan species. These solid clades are likely to derive from a single ancestral gene. We found four well supported clades (aLRT > 0.95), that we named clade I to IV, that contain a broad sampling of Bilaterian species, but no non-Bilaterian species. These are likely descendants of single pan-Bilaterian globin genes. In addition, one species rich clade containing both Bilaterian and non-Bilaterian genes (Porifera, Placozoa, Cnidaria) was found but with only a marginal aLRT support. We named this clade number V as we consider that it likely contains descendants of a single pan-metazoan gene. Last, distinct of these five pan-Bilaterian or pan-Metazoan clades, we found a well-supported clade containing only Spiralian genes (Mollusca, Annelida, Brachiopoda, Platyhelminthes), which we termed Sp-globins (Sp-gb). In addition to these multi-species clades, two other smaller clades are visible. One of them contains mostly globins of the Cnidarian *Nematostella vectensis* as well as a very derived globin of *Drosophila melanogaster* that might be there because of long branch attraction. The other clade contains only globins from the Nematode *Caenorhabditis elegans*. These mono-species clades presumably represent relatively recent gene radiations of a single divergent globin and remain unclassified. These eight well supported clades are connected by nodes with low support. Note that our arbitrary root chosen for representation in figure 3 (and additional file 5) is outside of these eight clades. We conducted bayesian phylogenetic analyses of the same sampling of genes and found good support for most clades (types I, II, III, IV, Sp-gb), but not for clade V (additional file 5).

**Figure 3:**
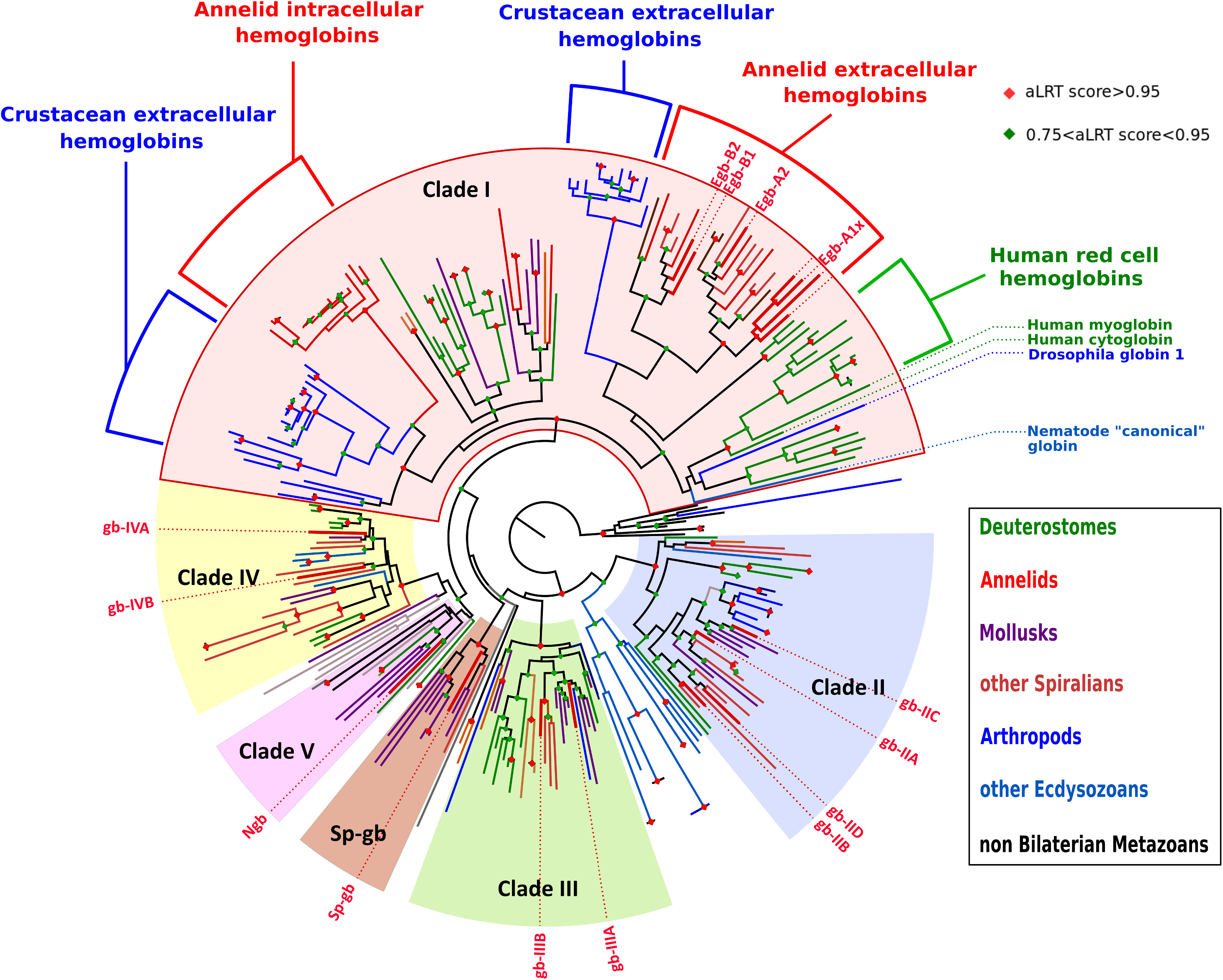
Maximum likelihood tree of Metazoan globin sequences. LG model of amino acid substitution was used and a SH-like test of a dataset of 272 sequences from 24 metazoan species. To improve readibility and facilitate annotations, a simplified tree version is used, where gene names do not appear and branches have been color-coded according to the clade of the metazoan tree in which they occur (colour code in the box on the right). The tree is arbitrarily rooted outside of the natural clades. Green and red diamonds indicate a number of nodes that are supported by aLRT values superior to 0.75 and 0.95, respectively. The tree shows 5 groups (clade I, clade II, clade III, clade IV, clade V), 4 of the 5 with well-supported nodes (clade I, clade II, clade III, clade IV), encompassing sequences of most Bilaterian phyla. An additional well-supported node groups together only Lophotrochozoan species sequences. *Platynereis* sequences are indicated by their names and thicker red branches.

We mapped on our ML tree the globins that are thought to be membrane bound because they harbor predicted dual N-terminal acylation signals (additional file 6; additional file 5, green rectangles) (4,21). None of the clade I globins is predicted to be membrane-bound. This is in contrast with the clade II, III, IV and Sp-gb, that all contain numerous predicted acylated globins. We retained as likely membrane-associated only those peptides that show strong predictions of both N-terminal myristoylation and palmytoylation. It is likely however that more globins of these clades are actually membrane-bound, for example by combining myristoylation with a basic motif. Based on the large proportion of predicted acylated peptides, there is a strong probability that the ancestral globins of clades II and IV, at least, were themselves linked to the membranes.

This analysis thus suggests that all Bilaterian globins derive from five ancestral genes that were present in *Urbilateria*, thus defining five clades of Bilaterian globins. Interestingly, *Platynereis* globins are distributed in all five clades plus the Spiralian-specific clade. In the genomes of non-Bilaterian Metazoans, a very variable number of globin genes are found. To analyse further the pre-Bilaterian evolution of globins, we performed a second ML analysis with an extended sampling of non Bilaterian Metazoans (sequences: additional file 7; tree: additional file 8), two Choanoflagellates and one Acoel. This extended analysis supports the presence of a single Clade V globin in the last common ancestor of Metazoans (additional file 6 and 8). The situation remains unclear in Cnidarians as Hydrozoans and one Ctenophore appear to have clade III-related sequences. Additional complete genomes of non Bilaterians and resolving the controversy of their phylogenetic relationships will be needed to understand fully the complement of globins in the last common ancestor of Metazoans.

All blood respiratory globins, or suspected to have a respiratory function, either intracellular (human) or extracellular (Annelids, *Daphnia*) are found in clade I in the ML tree (Figure 3). They also form species-specific clades inside clade I. All human red cell hemoglobin genes are found in a strongly supported clade with the human myoglobin and the human cytoglobin. All the *Daphnia* globin domains of their large family of didomain proteins are found in two separate clades, the N-domains and the C-domains, again suggesting a radiation from a single ancestral didomain-coding gene. All extracellular Annelid globins are found in one moderately supported clade, containing two globin "A" and "B" clades. These extracellular globins are present in four of the six Annelid species selected (*P. dumerilii*, *Arenicola marina*, *Lumbricus terrestris* and *Alvinella pompejana*), representing groups far apart in the Annelid tree. This grouping reinforce the interpretation that *P. dumerilii* putative extracellular globins have indeed a respiratory function, as this function is demonstrated by a number of work in the three other Annelid species (13,14,22). It also shows that the acquisition of a secretion signal and the gene A/B duplication that gave birth to these respiratory proteins must the have taken place early in Annelid evolution, possibly before the last common ancestor of all living Annelids. To strengthen this interpretation, we also searched two other Annelid genomes do not present any putative extracellular globins. One is the leech *Helobdella robusta* in which no clade I globins are detected. The other is *Capitella teleta* that present a large family of strongly related clade I intracellular globins, that are suspected to have a respiratory function (23). All the families of circulating respiratory globins present in distant groups in our analyses (vertebrate red cell globins, *Daphnia* extracellular globins, Annelid extracellular globins, *Capitella* red cell globins) thus correspond to independent event of gene recruitment and duplications, seemingly from a single ancestral clade I globin.

### Platynereis *extracellular globins are produced in specialized cells lining some vessels*

While our phylogenetic analysis suggests a circulating respiratory function for *Platynereis* extracellular globins, we wanted to obtain strong evidence of the linkage between extracellular globins and the development of vascular system of *Platynereis*. We particularly wanted to establish which cells are responsible for the production of *Platynereis* putative hemoglobin. To this end, we compared and combined two types of analyses: an expression analysis of globin genes by *in situ* hybridization *in toto* (WMISH) on a series of *Platynereis* developmental stages and a transmission electron microscopy study of juvenile stages to explore the vascular system development and characterize the globin producing cells.

*Platynereis* shows a complex succession of larval and juvenile stages, as well as a spectacular sexual metamorphosis, called epitoky (additional file 9). Specific RNA probes were produced for each globin gene. It must be noted however that the *Pdu-Egb-A1dα*, - *A1dβ* and *-A1dγ* genes have very similar nucleotide sequences, as well as being close chromosomal neighbors. The signal obtained with the *Pdu-Egb-A1dα* probe is probably a mix of the expressions of all three genes. None of the globin probes gave a significant signal on early larval stages (24 hpf trochophore, 48 hpf metatrochophore, 72 hpf nectochaete). The 10 intracellular globins gave no signal or non-specific signal in silk glands after long staining periods (not shown). Only six of the seven extracellular globin probes showed specific patterns. They all display expressions in the same cells, but not necessarily at the same time. Typical expression patterns for Egb-A2 are shown in Figure 4. In particular, we saw no expression of Egb-A1c in juvenile worms. The expression for this gene starts only in worms that are approaching sexual maturity and are about to enter epitoke metamorphosis. For all the most precocious extracellular globins, the expression starts when the worm is about 5 segments (not shown) in the posterior-most segment. In older juveniles, the expression is located along presumptive lateral vessels in the trunk (Figure 4A', red arrowheads). The expression shows a graded pattern along the anterior-posterior axis: stronger in the mid-body segments and absent in the rostral most and caudal most segments. This graded expression continues as the worms continue to develop. In bigger worms, the expression starts in the appendages of the mid-body along putative vessels that irrigate the part of the appendage that serves as a respiratory gill. The central most segments display expression in the parapodia when more rostral and caudal segments still display expression around lateral vessels in the trunk (Figure 4C', red arrowheads). There is seemingly a progressive migration of the hemoglobin-producing activity from the trunk to the parapodia and a more intense expression in the mid-body segments (Figure 4C; additional file 10). In worms engaged in sexual metamorphosis, we find a decreasing intensity of expression of all extracellular globins in parapodia (Figure 4E-F). In fully mature swarming worms, hemoglobin content is peaking but there is no mRNA of any blood globin left (Figure 4G).

**Figure 4:**
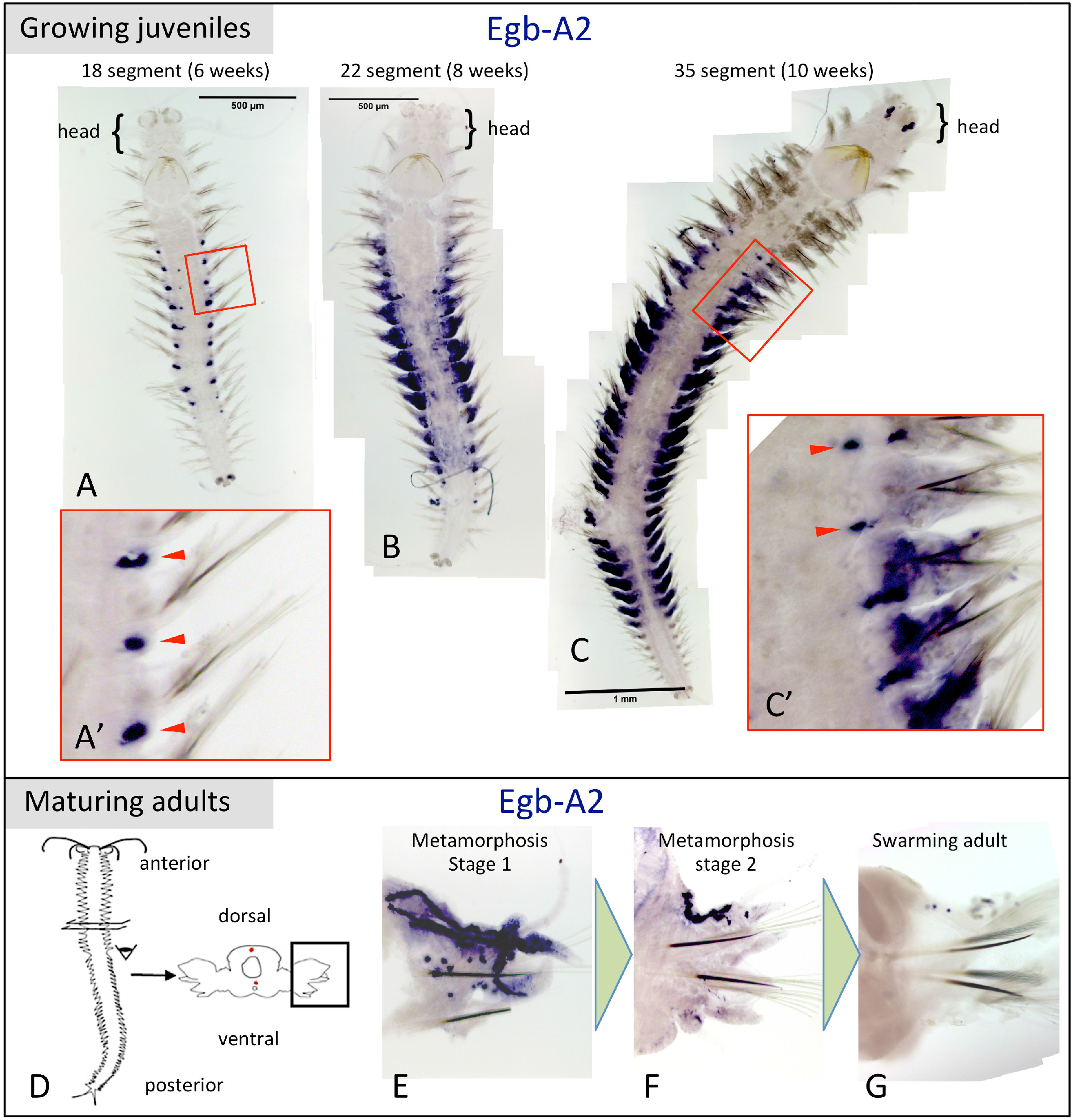
Expression patterns of *Pdu-Egb-A2,* a extracellular globin genes in *Platynereis dumerilii*. WMISH on three different juvenile stages. The expression patterns of all extracellular *P. dumerilii* globins are located in the same hemoglobin-producing cells of transverse trunk vessels and parapodial vessels. A - 18-segment juvenile show expression in transverse vessels in the trunk in a series of adjacent segments (red arrowheads). There is no expression in the rostral-most and caudal-most segments. A' - magnified view of three central segments. B and C - 22-segment juvenile and 35-segment juvenile show expression in the parapodia (segmental appendages) for central segments and expression in lateral vessels in a few segments more rostrally and caudally located. C' - magnified view of a few anterior segments of worm (C) showing some residual expression in transverse vessels (red arrowheads). D - drawing of a pre-mature worm explaining the dissection and mounting of parapodia. E-G - decreasing expression of *Egb-A2* in parapodia of worms undergoing sexual metamorphosis. Notice the delineation of vessels by hemoglobin-producing cells in the dorsal part of parapodium E.

Next, to understand how the *Platynereis* BVS develops and to localize the extracellular globin-producing cells and analyse their cytological properties, we observed stained semi-thin sections of worms at various developmental stages and made 3D reconstructions of their blood vessels (Figure 5A-J). We also observed ultra-thin sections of worms by electron transmission microscopy TM (Figure 5G-R). The *Platynereis* BVS is built progressively during development. The first signs of a BVS appear when a feeding juvenile with 3-4 segments has settled on the substrate around 15 days after fertilization, in the form of a pulsatile dorsal vessel and a non-contractile ventral vessel (Figure 5A-B). Then the juvenile starts to add segments at the posterior tip of the body and this sequential addition last during most of the benthic life of the animal (Figure 5C-F). New metameric BVS unit are put in place in each new segment added. Each metameric unit communicate with the contiguous segments by the pulsatile dorsal vessel, the quiescent ventral vessels and derivations of the lateral vessels and new vessels appear in the growing trunk, especially at the level of the segmental appendages, which serve both as legs and branchiae (gills).

**Figure 5:**
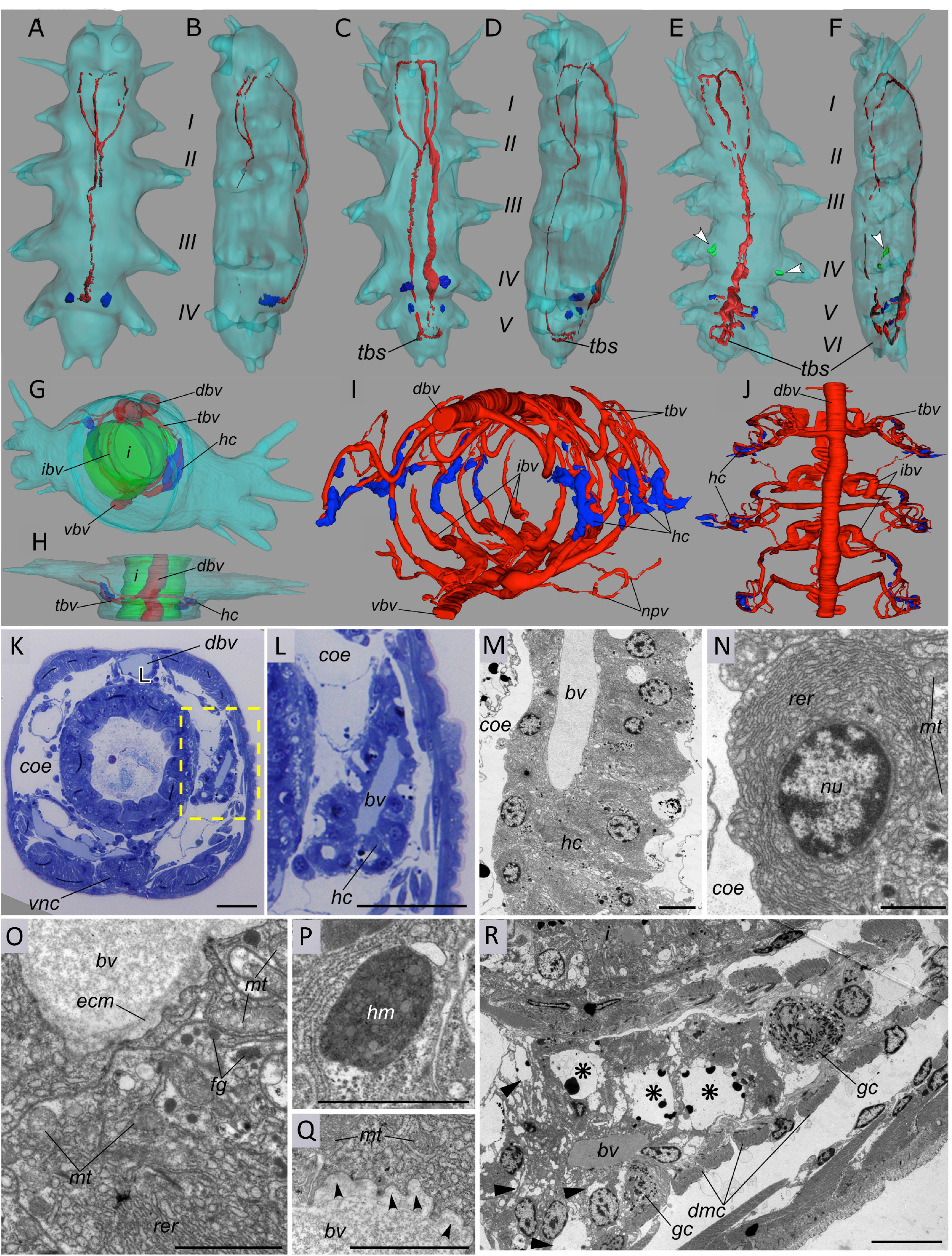
Developing vasculature and hemoglobin producing cells in *Platynereis* as revealed by semi-thin sections and TEM. A-J: 3D reconstruction of the blood vascular systems of early *P. dumerilii* juveniles. All reconstructions are made from stacks of semi-thin sections using Fiji TrackEM2 plugin. 4- (A-B), 5- (C-D), and 6- (E-F) segments juveniles of *Platynereis dumerilii*. (A-C-E) Ventral views. (B-D-F) Lateral left views,. tbv: transversal blood vessels. G, H: single segment of a 25-segment juvenile worm, carrying a pair of appendages (parapodia) in anterior view and dorsal view, respectively. The outside body wall is lined in light blue. The intestine endoderm is in green. I, J: Vascular system in three contiguous segments of a 50-segment sub-adult worm. For the sake of clarity, the dense intestinal blood network (or gut blood sinus) is not shown. *dbv* – dorsal blood vessel, *hc* – hemoglobin-producing cells (blue), arrowheads – putative degenerative hemoglobin-producing cells (green), *i* – intestine, *ibv* – intestinal blood vessels, *npv* – nephridial vessels, *tbv* – transversal blood vessels, *vbv* – ventral blood vessel, *tbs* – transverse blood sinus (pygidium), all vessel lumens are in red. K-R: Ultrastructure of *P. dumerilii* hemoglobin-producing cells in a juvenile worm, 25-30 segments. K: transversal semithin section, showing the position of hemoglobin-producing cells around the transversal blood vessel. L: magnified view of the dissected transversal vessel surrounded by hemoglobin-producing cells (rectangle selection in K). L: Transmission electron micrograph of the overall hemoglobin-producing tissue organization. M-R: Transmission electron micrographs of different parts of the hemoglobin-producing cells. M: Apical part of the cell, containing nucleus (*nu*) and rough endoplasmic reticulum (*rer*). N: Basal part of the cell with mitochondria (*mt*), rough endoplasmic reticulum (*rer*), and ferritin granules (*fg*). P: closer view of a ferritin granule (*fg*). Q: Invaginations of the basal membrane (arrowheads) indicating active exocytosis process. R: Transmission electron micrographs of the degenerating hemoglobin-producing cells in sub-adult *P. dumerilii* (about 50 segments); cross-section through the transversal blood vessel at the level of hemoglobin-producing cells. Notice the cytoplasm, filled by electron-lucent vesicles (*arrowheads*), the large electron-lucent vesicle (*asterisk*) and the granulocytes filled with electron-dense and electron-lucent granules. *bv* – blood vessel lumen, *dmc* – muscular cells of the dissepiment, *gc* – granulocyte, *nu* – cell nucleus, *coe* – coelomic cavity, *dbv* – dorsal blood vessel, *ecm* – extracellular matrix (vascular lamina), *hm* – hematin granules, *hc* – hemoglobin-producing cells, *mt* – mitochondria, *nu* – cell nucleus, *rer* – rough endoplasmic reticulum, *vnc* – ventral nerve cord. Scale bars: K-L: 50 μm, M: 10 μm, N: 4 μm, O-Q: 2 μm, R: 10 μm.

The blood becomes visibly red only when the worm reaches a certain size, around 10-15 segments. This suggests that blood hemoglobin producing cells become active progressively. One stage corresponds to juvenile worms with around 25-30 segments (Figure 5G-H). In these worms, the BVS metameric unit is still fairly simple with dorsal and ventral vessels and a couple of lateral vessels (Figure 5G-H, red tracks). The lateral appendage vasculature is still little developed. At this stage, sheaths of characteristic putative hemoglobin-producing cells (hc, in blue) are already present around a pair of transverse blood vessels (tbv, Figure 5G-H). The other stage corresponds to pre-mature worms with around fifty segments. These worms have much more developed lateral appendage network with lateral vessels colonizing the gill-like parapodia (Figure 5I-J, red; movie in additional file 11). In the mid body segments of these worms, the putative hemoglobin-producing cells sheaths are still visible around the transverse vessels (tbv) but new hemoglobin-producing cells appear around parapodial vessels, more densely on the dorsal side.

Putative hemoglobin-producing cells semi-thin and TM images are illustrated in Figure 5K-R. On semi-thin sections, they appear as thick sheaths of cells surrounding the lateral vessel lumen (Figure 5K-L). The hemoglobin-producing cells are organized in a simple epithelium with basal sides facing the vessel lumen and apical sides facing the coelomic cavity (Figure 5M). A clear basal lamina delineates the vessel lumen on the basal side (Figure 5O). Hemoglobin-producing cells are therefore meso-epithelial cells in clear continuity with the meso-epithelium that surrounds the coelomic pockets. They show all the characteristics of an intense protein synthesis and secretion activity (Figure 5C-G). The cytoplasm is mainly filled with a dense rough endoplasmic reticulum (Figure 5N-O). Electron-dense bodies are visible in the cytoplasm of hemoglobin-producing cells and are interpreted as ferritin granules (Figure 5P). Last, secretion vesicles are clearly identified on the basal side (Figure 5Q), indicating intense exocytosis process. The vesicles contain a uniformly granulated matrix identical to the aspect of the blood. These small granules are thought to represent the particles of the giant hexagonal bilayer of erythrocruorin, characteristic in many Annelid species. These hemoglobin-producing cells are remarkably similar to the previous descriptions of hemoglobin secreting cells in the heart-body (an organ inside the dorsal vessel) of several Sedentarian Annelids (24). Small juvenile worms with 4-6 segments already have hemoglobin-producing cells, visible in the posterior most segments as small groups of cells (blue) (Figure 5A-F).

In juveniles of various stages, we also identify secretory cells with clear signs of apoptotic degenerescence (figure 5R). We believe they represent dying hemoglobin-producing cells. Their location is correlated with the globin-production shift we detected in the WMISH analysis. We find degenerating cells around the transverse vessels in the mid-body segments at the time when globin production shifts from the trunk to the appendages. We find these degenerative cells as early as in small juvenile with 6 segments (Figure 5E-F, in the forlast posterior segment, green).

### Evidence for globin isotype switching during the life cycle

In addition to the spatial pattern evolution, our WMISH analyses suggest that different extracellular globin isotypes might be expressed at different times. To establish strong evidence, we turned to quantitative PCR analysis.

The stages for qPCR were divided as follows: 48hpf larvae, 6 weeks larvae, 50 segments larvae, sexual metamorphosis stage I, sexual metamorphosis stage II, mature swimming worms. We designed specific primer pairs for each Egb gene, but again it is expected that primers for *Pdu-Egb-A1dα* are going to capture expressions of the two closely related genes *-A1dβ* and *-A1dγ* as well. The relative expression levels of the extracellular genes with respect to reference genes show important variations from one biological triplicate to the other (additional file 12). Different individuals of the same apparent stage can display substantially different levels of globin mRNA. We have no clear explanation for this important variability but globin expression at the mRNA level is known to be highly responsive to physiological conditions, in particular the amount of O_2_ available in the environment (25). Notwithstanding, the general trend is that blood production picks up at 6 weeks, culminates between 50 segments and the beginning of the sexual metamorphosis (75-80 segments) and collapses rapidly after, ending by the time the worm starts swimming, which will be followed by reproduction and death (Figure 6). Individual gene expressions, despite variability, show trends that confirm our WMISH observations for some genes. The globins A2, B1 and B2 (Figure 6) are expressed at levels that follow the general tendency described above and at higher levels than the four A1 globins. The four paralogous A1 globins (Figure 6), meanwhile, are expressed at different levels. While A1a and "A1d" follow the general tendency, A1c is expressed significantly only in sexually maturing animals maybe representing an adult-specific erythrocruorin sub-unit. In contradiction with at least some of our WMISH results, A1b was detected only at very low levels in all biological replicates.

**Figure 6:**
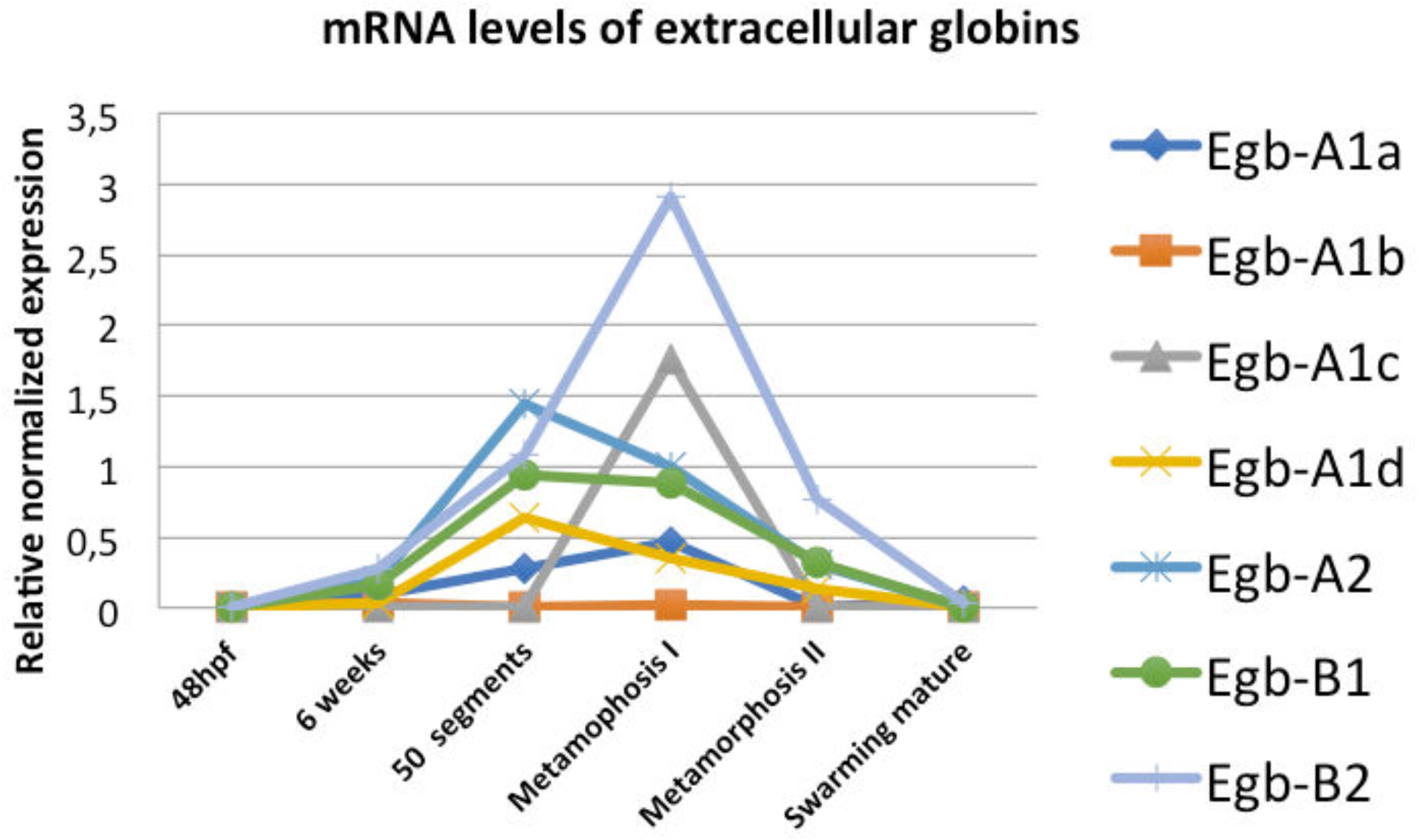
Relative abundance of mRNA of most *P. dumerilii* extracellular globin genes, detected by real-time qPCR, at different life cycle stages. Metamorphosis stage 1 is defined according to the following criteria: eyes starting to bulge, transforming parapodia. Metamorphosis stage 2: eyes grown to maximal surface, parapodia fully transformed, but the animal is not displaying active swimming behaviour yet. Reference genes are *rps9* and *sams*. For stages 50 segments, stage I pre-matures, stage II pre-matures and swarming worms, the data points correspond to the median value of biological triplicates.

## Discussion

We report the first exhaustive screening for globin genes in an Annelid and more generally in a Protostome animal. We performed a broad phylogenetic study of the metazoan globins based on complete genome sequences. This study helps in understanding the evolution of the globin superfamily in Metazoans. Combined with the existing knowledge, it gives some hints in the evolution of the red blood itself. Hemoglobins evolved several times but from a unique intracellular stem globin, which appears to be still present in most Bilaterians, the clade I stem globin or cytoglobin as it is termed in Mammalians.

### A phylogenetic nomenclature for Metazoan globins

Our study of the phylogenetic relationships of metazoan globins fills an important gap in the literature. Many studies have actually been published on the phylogeny of globins starting with the first molecular tree (3). To cite only some of the recent works, there has been studies on the evolution of globins at the level of the tree of life (1,26), and at the level of the eukaryotic tree (27). These studies emphasize the universality of the globin fold motif present since the last common ancestor of all living cells (LUCA, last universal common ancestor) and the likely functions of these proteins as enzymes or sensors. Many other studies have analysed globin evolution in specific metazoan groups such as Annelids (28,29), Deuterostomes (6), Echinoderms (20), Pancrustaceans (30), Insects (31), Cephalochordates (32), Chordates (33), Agnathans (34,35). This proliferation of studies illustrates the extraordinary diversification of globin genes in many of the metazoan groups.

The previous most comprehensive studies (4,5,7) have exemplified the existence of large conserved sub-families of non-blood globins in Metazoans and have started to show the complexity of animal globin evolution. One of these sub-families, the neuroglobins (Ngb) are hexacoordinate intracellular globins first discovered in the mammalian nervous system and whose function remain poorly understood (8,36,37). Neuroglobin orthologues have been discovered in a number of animal groups other than Vertebrates as well as in Choanoflagellates and has been proposed as an ancestral Metazoan globin from which all the other animal globins could be derived by gene duplication (7). The other proposed sub-family has been called “Globin-X” (9). Globin-X (and Globin-X-like) proteins are widespread among Metazoans. They present very often one or two acylation sites at the N-terminus, either myristoylation or palmitoylation, making them membrane-bound proteins (4). The functions of globin-X proteins are not known, although a role in protecting the cell from reactive oxygen species (ROS) is suggested. Blank and Burmester’s work (2012) suggest that the blood globins of Vertebrates may derive from ancient membrane-bound globins, via the loss of acylation sites, opening, among other things, the possibility to form multimers.

Despite bringing forward interesting conceptions, previous works on Metazoan globins all suffer from biased sampling. The sampling of globins of Blank and Burmester (2012) was made by searching NCBI Genbank with terms related to globins, while the same database was screened with a Ngb probe in Lechauve et al (2013). In this work, we have screened complete genome sequences of a representative sampling of metazoans, using a concatemer sequence probe representing the diversity of human and *Platynereis* sequences. We are thus confident that our samplings of globins in each genome are very close to exhaustivity. In trees based on maximum-likelihood and bayesian algorithms, a number of robust nodes emerged that correspond to globins present in a large sampling of Bilaterian species, thus likely to represent ancestral Bilaterian genes.

The existing classification of globins is mostly Mammalian and function oriented. This is of course quite relevant in certain context. In this article, we want to give a different viewpoint by providing a classification that is entirely based on gene phylogeny at the Metazoan level, exactly as it exists for other gene families. Previous publications have tended to develop additional nomenclature by keeping strong mammal-and hemoglobin-centered (hence an evolutionary derived function) points of view. Globins evolved in functional contexts that were entirely different from the eventual evolution of a blood function in a few animal groups and this is what our phylogenetic nomenclature reflects.

The five ancestral groups of globin molecules we identified (Figure 7A) are the following:

- Clade I: this includes the totality of circulating blood globins, either extracellular or carried by red cells, found in metazoans. In addition, this group also includes many non-circulating globins. These additional non-blood clade I globins are present in animals with hemoglobin blood, with a non-hemoglobin blood or with no blood at all. The large majority of Bilaterian taxa are represented in this group but no globin of non-Bilaterian taxa (Cnidarians or Sponges) is present, indicating that this clade originated in an ancestor of Bilaterians. Globin proteins can display two different chemistries, depending on whether the heme is attached to the globin by one or two histidine side chains, respectively referred to as “pentacoordinate” or “hexacoordinate” (38). The Vertebrate hemoglobins and myoglobins, responsible for conveying and storing oxygen are pentacoordinate. However most globins of clade I are hexacoordinate suggesting that the Vertebrate blood globins have made the transition from hexacoordination to pentacoordination.
- Clade II: this clade contains multiple globins from all Bilaterian superphyla (Deuterostomes, Ecdysozoans, Spiralians). It has been called formerly « globinX-like » (4). Clade II globins are not identified in non-Bilaterian taxa, indicating that this clade originated in an ancestor of Bilaterians. No clade II globin is present in Vertebrates (although it is present in Cephalochordates), indicating that a clade II globin was present in the common ancestor of Chordates and lost secondarily in Vertebrates.
- Clade III: this clade contains globins from all Bilaterian superphyla. It corresponds to « globinX » (4). Clade III globins are not identified in non-Bilaterian taxa in our main sampling, but the presence of Clade III-like genes in Hydrozoans and a Ctenophore indicates that this clade may have originated in an ancestor of Eumetazoans. Our sampling contains only one Vertebrate species, human, in which clade III globins are not present. It is known from other studies that clade III globins are present in non-Amniote Vertebrates, but absent from Amniotes, indicating secondary loss.
- Clade IV: this clade contains globins from all Bilaterian superphyla, although it is absent from Chordates and also from all Ecdysozoans except Priapulida. This clade has gone undetected in previous studies. Clade IV globins are not identified in non-Bilaterian taxa, indicating that this clade originated in an ancestor of Bilaterians.
- Clade V: this clade is the least statistically supported of all. It contains however globins from a broad variety of Metazoans, including Sponges and Cnidarians, as well as Choanoflagellates, one of the closest Eukaryotic sister group of Metazoans (additional files and 8). This is the only globin type whose existence is clearly suggested in the last common ancestor of Metazoans. Paradoxically, it is also the most "dispensable" globin. It has disappeared from the genomes of the majority of the species in our sampling (especially all Ecdysozoans). Clade V corresponds to some of the « neuroglobins » identified in previous studies but this is however a much narrower group than the « neuroglobin-like » proteins previously described (7). These authors included many globins in a broad « neuroglobin-like » class, which fall in clade II and III in our study.
- In addition, a well supported clade contains only globins from Spiralian species (Sp-gb). It may have appeared by a gene duplication of any of the other globin clades.

**Figure 7:**
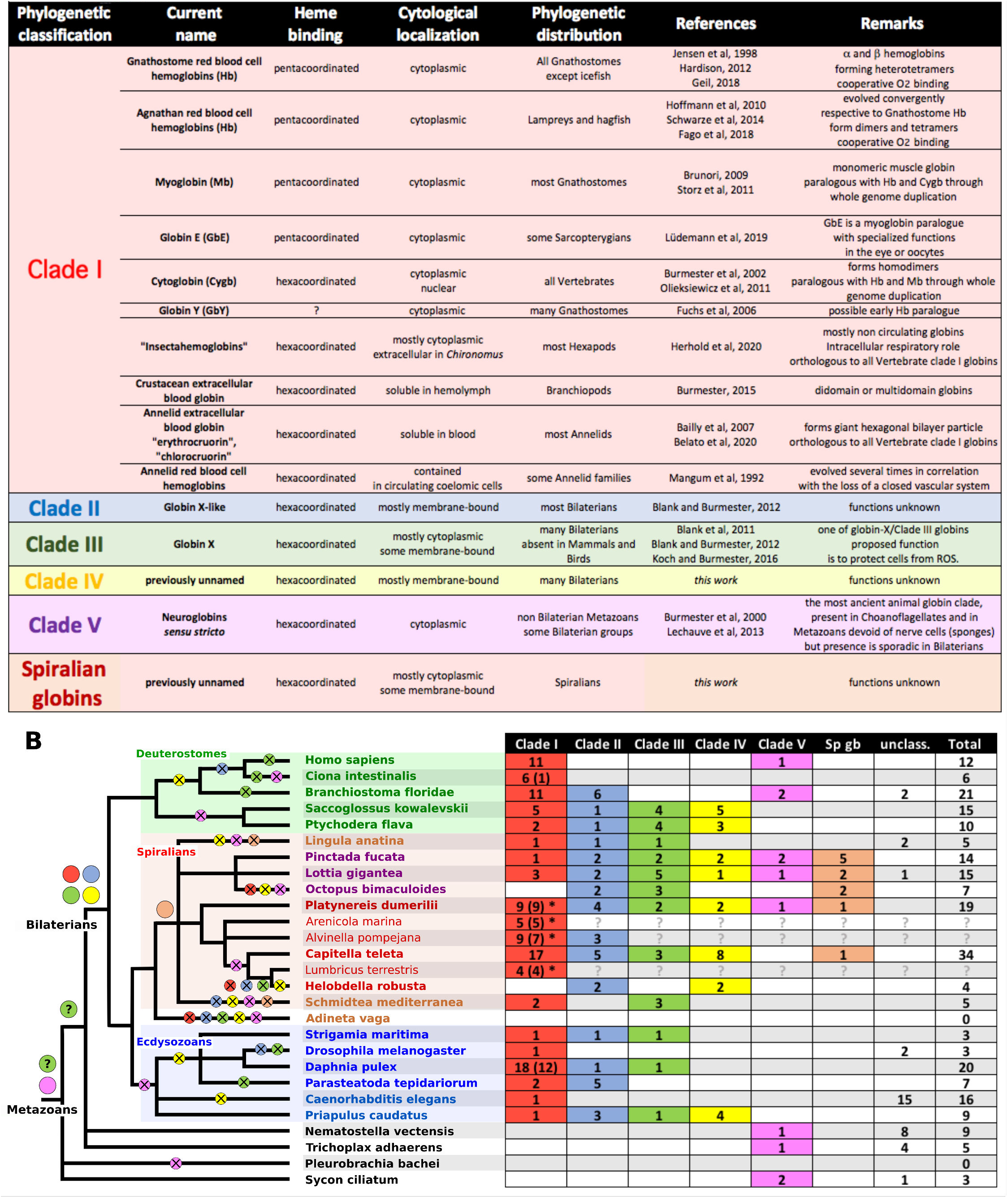
Phylogenetic classification of Metazoan globins and evolutionary scenario for their evolution. A - Table of the clades suggested in this study. The current name column indicates the previous denominations that have been used in previous literature. B - Table of the number of genes of each clade present in the species genome we sampled. For the clade I, the number of extracellular globins is indicated between parentheses. An asterisk shows all the species in which hexagonal bilayer linker proteins have been detected. On the right, the animal tree shows the inferences of gain and loss of globin gene clades. Colored disks indicate the gain of a new gene clade by duplication from existing ancestral genes while crossed disks illustrate the numerous secondary losses of globin clades. Interrogation marks indicate missing data because the whole genome is not available for the relevant species.

### The animal globin superfamily has an ancient history independent from the history of blood

The existence of these clades clearly indicates that at least five globin genes existed in *Urbilateria*. We call these five hypothetical molecules the “Bilaterian stem-globins”. Few globin sequences fall outside of these five well-defined clades (Figure 7B). These additional globins fall into two cases. In one case, these sequences may derive from the stem globins but could have evolved rapidly so as to blur sequence similarities. This is what we would suppose for the large clade of *C. elegans* globins found alongside a single clade I globin. The second case concerns the clade of sea anemone globins, which includes almost all the sea anemone globins in our dataset (except from a single clade V globin). Most of the members of this group are membrane-bound, suggesting that it could be related to any of the Bilaterian stem globins that were potentially membrane-bound (clade II, III and IV). Alternatively, the sea anemone membrane-bound globins may derive from an ancestral Eumetazoan membrane-bound globin that would also have given birth to Bilaterian clade II, III and IV. Reconstituting Metazoan globin history prior to Bilaterian diversification is thus at this stage still tentative because of the small sampling of species.

The deep relationship between Bilaterian stem globins is not solved by our trees (that are arbitrarily rooted in any case). The idea that Bilaterian blood globins (and all other clade I globins as defined by our trees) may derive from molecules that were initially membrane-bound (4) is thus not corroborated by our study. It is in our view equally possible that clade I derives from an ancestral duplication of the cytoplamic clade V, while clade II, III and IV would have acquired membrane tethering independently.

What were the functions of these numerous non-blood globins derived from the Urbilaterian stem-globins? Here again, very few functional studies have been performed and we can only speculate. Burmester and Hankeln (2009) have reviewed the possibilities for the special case of neuroglobins (clade V), but the same possibilities seem to exist for the other clade of non-blood globins (clade I to IV). These intracellular globins, cytoplasmic or membrane-bound, or in some cases located in the nucleus (39), can display a “classic” function, similar to the Vertebrate myoglobin (Mb) in storing O_2_ in hypoxic conditions or facilitating the delivery of O_2_ to the mitochondria. New functions have also emerged in the literature, such as the regulation of reactive oxygen species (ROS) (40) or reactive nitrogen species (RNS) that can be deleterious to the cell. It has been also been speculated that they could function as O_2_ or redox sensors by interacting with other proteins such as G proteins (41) or cytochromes to transmit a signal.

The five Urbilaterian stem globins have undergone remarkably dissimilar fates in different Bilaterian groups (Figure 7B). They have all been conserved in the Annelid *Platynereis dumerilii* and the Mollusks *Pinctada* (Bivalvia) and *Lottia* (Snail). In other Bilaterian groups, one or more of the ancestral clades have been lost. Amniotes have kept only clade I and clade V globins. The Urochordate *Ciona* has only clade I globins left but six clade I paralogues are present, maybe compensating for the loss of the other types. The most extreme case was however the Rotifer *Adineta,* in which we could not identify any globin-related gene. This species has thus lost all five ancestral globins.

### A novel scenario for the evolution of blood globins

Oxyphoric circulatory globins have evolved several times during the course of Metazoan evolution. All circulatory multigenic families in our dataset derive from a likely unique clade I globin, possibly after recruitment for the circulatory function and several rounds of gene duplications. These multigenic families encompass for example the human hemoglobin family as well as the extracellular globin families of Annelids and *Daphnia*.

In the case of the hemoglobins of Vertebrate erythrocytes (RBC), heterotetramers comprising alternate globin isotypes circulate depending on the development stage. Adult human hemoglobins form heterotetramers comprising sub-units of two isotypes, α and β, coded by paralogous genes (42). During the course of human embryogenesis and foetal development, different isotypes of hemoglobins are produced (δ, γ, ε, ζ), coded by other paralogous genes located in two clusters of α and β-related paralogues. The crucial emerging property of the hemoglobin tetramer is cooperativity: the binding of an O_2_ molecule to one of the sub-unit induces a conformational change in the tetramer that makes the binding of O_2_ easier to the three remaining sub-units.

Elsewhere among Metazoans, other gene radiations correlated with hemoglobin-based blood evolution have occurred. Three of the concerned groups are sampled in our tree: Four species representing a wide diversity of Annelids, the special case of the Annelid *Capitella teleta*, the Pancrustacean *Daphnia*.

In Annelids, our phylogenetic analysis is in accordance with the scenario that early gene duplications, probably in the last common ancestor of modern Annelids, produced the initial two families (A, B) encompassing the four extracellular globin sub-families A1, A2, B1, B2 that form the hexagonal bilayer hemoglobin assemblage. A recent work based on a massive sampling of Annelid species transcriptomes recovered a variable number of extracellular globins, ranging from 1 to 12 depending on the species (29,43). Gene trees made in this later work suggest that, while the early gene duplication A/B indeed occurred prior to Annelid diversification, the sub-families A1, A2, B1 and B2 are not recognizable at the scale of the Annelid tree. This is in accordance with what we obtain at the much smaller scale of our Annelid sampling. This fact might be interpreted as evidence that the Annelid last common ancestor has only two clade I paralogous globin genes, A and B, and that additional gene duplications occurred independently in many Annelid lineages. Alternatively, four paralogous genes (A1, A2, B1, B2) may have existed early on in Annelid history, accounting for the hexagonal bilayer hemoglobins found in divergent Annelid groups, but in this case the gene sequences are not informative enough to reconstitute early duplication steps. Our work differs however in a striking way from previous results (43). These authors identify a number of extracellular globins related to the Annelid A and B extracellular globins from a large and diversified subset of Bilaterian species including Echinoderms, Hemichordates, Brachiopods, Nemertines, Mollusks and Priapulids. We find no evidence for the existence of globins related to the Annelid extracellular globins (i.e. hexagonal bilayer globins) in Bilaterian complete genomes, other than Annelids. We argue for a thorough reconsideration of transcriptome screens (43), as many protein globin sequences these authors found in non-Annelid Bilaterians are strikingly similar if not identical to individual Annelid sequences (as illustrated in their Figure 4). This is suggesting multiple tissue or DNA contaminations between species transcriptomes. Despite *Platynereis* blood hemoglobin having never been directly characterized by spectrometric tools, our EM and sequences similarities show that *Platynereis* must exhibit the giant assembly. In *Platynereis*, additional duplications have affected the A1 gene, making several paralogue sub-units. These sub-units, according to our *in situ* hybridization and qPCR data, may be used in producing alternative composition of circulating hemoglobin that would be used for physiological adaptations as the worm develops, grows and metamorphoses.

The clade I stem-globin evolved into a family of intracellular globins in some Annelid species. In the case of the Sedentarian worm *Capitella*, which does not have blood vessels, the hemocoel contains many red cells with intracellular globins with oxygen binding properties (11). We have identified 17 intracellular globin paralogues, all closely related, in the genome of *Capitella* SpI and no extracellular globins. It appears as a case where the ancestral Annelid extracellular globins were lost secondarily and replaced by RBC globins.

Although hemocyanin is the main respiratory pigment in Pancrustaceans (36), Branchiopods such as *Daphnia*, a few Malacostracans and a few Insects (such as *Chironomus* larvae) rely on hemoglobin for respiration*. Daphnia* hemoglobins have been thoroughly studied (44), as a model of adaptation to the environment by the extensive use of alternative globin isotypes. *Daphnia* possesses a large family of extracellular two-domains globin-coding genes. It is proposed that the hemolymph of *Daphnia* contains a crustacean hemoglobin made of 16 identical globin peptides with two hemes each (45). For our phylogenetic analysis, we treated the two globin fold sequences independently, resulting in trees indicating that the multigenic family of *Daphnia* globins has emerged from the tandem duplications of an ancestral didomain globin gene.

However, globin gene radiations are not necessarily linked to recruitment for a function in the blood. This is exemplified by Urochordates and Cephalochordates. The ascidian *Ciona intestinalis* possesses 6 clade I globins, all closely related. The cephalochordate *Branchiostoma* has eight clade I globins found in two clusters in this analysis. Both species have nevertheless a colorless blood and to date, no functional study has shown either oxygen binding properties or their presence in the circulatory system.

All clade I globins likely derive from a single ancestral clade I sequence. This ancestral clade I globin was duplicated several times in different clades independently to give small families of blood respiratory proteins. This raises two questions: What was the ancestral function(s) of this clade I globin that made it so well pre-adapted for a role in blood function and is there a gene, in current Bilaterian species, that still carries this ancestral function(s)?

We can only speculate at this point because little is known on the function of the non-circulating clade I globins in the vast majority of Bilaterian species. But the existing knowledge can at least help defining the direction of future research. Part of the answer may come from Vertebrates that possess both circulatory and tissue-expressed clade I globins. The “tissue” clade I globins in Vertebrates are myoglobins and cytoglobins. Myoglobins are monomeric clade I globins present in all Vertebrates. Their function is well known and very specific: they are expressed in muscles and are capable of storing large quantities of O_2_ necessary for sustained muscular effort. They also facilitate intracellular transport of O_2_ (46). Cytoglobins were first identified as distinct intracellular globins in Amniotes (18,47). They harbour a structural difference with hemoglobins (and myoglobin) in having a hexacoordinated iron, rather than pentacoordinated. They form homodimers, also by contrast to the former. Phylogenetically, they are firmly identified as clade I globins in our study. Contrary to myoglobins, they are expressed in a large range of tissues and organs, with maybe higher expressions in cells producing large amount of cell matrix such as fibroblasts and chondroblasts (48). Information on the function (or functions) of cytoglobins remains scarce to this date. It is known that they bind dioxygen, carbon monoxide but also nitric oxide with high affinity (49). It has thus been proposed that cytoglobins could play multiple roles in intracellular homeostasis. The first suggestion is of course related to dioxygen binding including roles in oxygen buffering, sensing, transport and storage. The activity of cytoglobin as nitric oxide deoxygenase, a very ancient function of globin proteins in eukaryotes, is also proposed to play an important role in the biology of the cell. Last, oxidative stress is another circumstance in which cytoglobins may play a role (49).

Remarkably, molecular phylogenies have demonstrated that large families of oxyphoric circulatory hemoglobins have evolved by gene duplications two times in the Vertebrates: once in the Gnathostomes and once in Agnathans (34,50). Interestingly, in these molecular trees, the cytoglobins of Gnathostomes and Agnathans are found as sister groups to both their large radiations of circulatory globins, suggesting that cytoglobin existed prior to the origin of red cell globins in Vertebrates. Cytoglobins evolve conservatively (33), suggesting that they keep important functions. There are good reasons to propose that cytoglobins (rather than the more specialized myoglobin) may be close to a clade I globin with ancestral functions, as was suggested before (4).

The fruitfly *Drosophila*, like other insects, has a tracheal respiration and no circulatory globins. Yet, genome screens have revealed three globin genes in the *Drosophila* genome (51). Two of these globins possess derived sequences. But the third one (Dme_524369; “Glob1” in Burmester et al, 2006) is found solidly clustered with clade I globins in our tree (Figure 3, additional file 5). This globin is hexacoordinated and may represent a derived cytoglobin-like molecule. It is found expressed in many tissues in the developing embryo and larva (52) but prominently in the tracheal cells. Gene knock-downs (52,53) lead to reduced survival of flies under hypoxic conditions. This suggests a role in intracellular O_2_ homeostasis.

Both cytoglobins and *Drosophila Glob1* may thus be close to the functions of a clade I stem globin. It has to be noted from the ML tree (figure 3) that several species (Arthropods and Mollusks) have a single clade I globin that may represent the unduplicated descendants of the stem globin. Few species have completely lost the clade I (only the Annelid *Helobdella* and the Cephalopod *Octopus*). This may reflect the initial functional importance of clade I stem globin.

Why have multiple gene duplications affected the clade I stem globin each time it has been recruited for a blood function? As we have mentioned earlier, cooperativity is an important functional mechanism for a multimeric globin whose role is to store large quantity of O_2_ in the blood, yet be able to deliver it on demand to all tissues in need. It may be best developed in a heteromultimeric assemblage and thus requires gene duplication and rapid divergence of the O_2_ storing protein. This idea has received crucial support from recent biochemical studies using synthetic ancestral proteins inferred from the diversity of extant Vertebrate globins (54). The second factor is the developmental switch of globins. This is well illustrated by the lamprey and the Gnathostome cases. The lamprey *Petromyzon* possesses 18 hemoglobin genes that are expressed differentially at three different developmental stages, embryo, larva and adult (34). These developmental switches are similar to those found in Gnathostomes. Yet, all the Agnathan circulatory globins are found in a single monophyletic group completely distinct from the radiation of Gnathostome hemoglobins (50). The lamprey globins family was convergently recruited for the very same kind of developmental as seen in Gnathostomes, We may see to a more limited extent the same developmental switch in *Platynereis*, with different A1 isotypes specialized for the juvenile and the adult.

### The hemoglobin-producing cells: a specialized type of blood-making cells

Little was known on the nature of the cells that produce and secrete respiratory pigments, outside of Vertebrates.

We demonstrate in this work the existence of a specialized category of vessel-lining cells in *Platynereis*, that produces the blood globins, in other words the hemoglobin-producing cells. We analysed the development of these cells during the life cycle of *Platynereis*. They first appear in the posterior most segments in a five-segment worm. The youngest feeding juvenile worms with 3 or 4 segments probably do not have hemoglobin-producing cells and therefore no hemoglobin in their blood. These are minute worms (less than 500 μm long) that probably perform gas exchanges with the seawater by simple diffusion through their tissue. In intermediate size juvenile worms (25-40 segments, less than one cm long), hemoglobin-producing cells form sheaths around transverse trunk vessels, in most segments of the worm. As the worm grows, the circulatory system structure becomes more complex and a rich network of vessels and capillaries develop inside the parapodia, the worm segmental appendages. These parapodia take on the role of gills. Hemoglobin-producing cells develop around most vessels inside the parapodia. We currently do not know whether these are the cells of the wall of existing vessels that are differentiating *in situ* to become hemoglobin-producing cells, or whether hemoglobin-producing cells colonize the parapodia by emigrating from another location. We also show that at this stage (worms larger than 40 segments), hemoglobin-producing cells within the trunk degenerate and are presumably digested by eleocytes. Worms accumulate more red blood as they get close to sexual metamorphosis. When they have a maximal quantity of blood and start metamorphosing into swarming epitokes, the hemoglobin-producing cells stop progressively their production. It is possible that these cells degenerate but we have not demonstrated this phenomenon in metamorphosing worms.

To our knowledge, this is the first time the full cycle of these hemoglobin-producing cells is described in any Annelid species and this is also the first time they are found associated with the parapodia serving as gills. Hemoglobin-producing cells with very similar cytological properties have been described in other Annelid species. They have been called perivasal cells because of their position around the lumen of vessels or extravasal cells, when they are located in the mesodermal peritoneum covering the gut in earthworms (55), Siboglinids (56) or *Arenicola* (57). In some Sedentarian Annelids such as *Amphitrite* (58), a specialized organ, the heart-body, produces the hemoglobins. The heart-body is a solid mass of tissue that grows inside the lumen of the anterior dorsal vessel (59). There is thus a remarkable plasticity in the location of this hemoglobin-producing tissue in Annelids that will be worth studying further. These studies reveal an unsuspected variability in the localisation of these cells in Annelids.

How do these hemoglobin-producing cells relate to the hematopoiesis processes? It should be noted that, in *Platynereis* and in Annelids in general, hemoglobin-producing cells are not circulating, contrary to Vertebrates RBC. They are part of a mesodermal epithelium that is in contact with the coelomic cavity on the apical side and with the blood vessel lumen on the basal side. They are functionally polarized cells that secrete hemoglobin only on the basal side toward the vessel lumen. Besides hemoglobin-producing cells, *Platynereis* (and Annelids in general) have several classes of coelomocytes/hemocytes that are freely floating in the coelomic cavity and can cross the mesoepithelium to populate the blood. Can hemoglobin-producing cells be considered as cells produced by an extended hematopoietic process? This is not only a question of convention, when considering the diversity of hemoglobin-producing cells in Metazoans.

Is there a phylogenetic connection between the different Metazoan hemoglobin-producing cells or more generally speaking between the different Metazoan respiratory pigment-producing cells (hemoglobin, hemocyanin, hemerythrin)? In the various Bilaterian groups that possess either a hemocoel or coelom/blood compartments, hemoglobin-producing cells can be either free floating or static. In Vertebrates, intracellular globins are massively produced and stored in red blood cells. Accumulating respiratory pigments in circulating cells is also the solution retained in a number of other groups. Within Annelids, the bloodworm *Glycera* have red cells filled with monomeric and polymeric hemoglobins, circulating in the coelom as the BVS is much reduced in these worms (60). Capitellids have also a reduced BVS and red cells. Also in Annelids, Sipunculidae have pink cells filled with hemerythrin, illustrating the evolutionary pigment swapping which has occurred in several groups. Data on the existence and location of pigment producing cells in other protostome invertebrates are still patchy. In the marine Chelicerate *Limulus*, whose remarkable blue blood has been exploited for preparing bacterial endotoxin tests, the hemocyanin is secreted by cells called cyanoblasts. Cyanoblasts are found floating in the hemocoel and burst to liberate their hemocyanin content (61). The tissue of origin of these cyanoblasts is unknown. In Gastropod Mollusks, cells responsible for hemocyanin production have been identified as the pore cells or rhogocytes (62), abundant in the connective tissue. In pulmonate snails that have hemoglobin instead of hemocyanin, it has been shown that these rhogocytes have switched to hemoglobin production (63). One can hope that in the future single-cell transcriptomics as well as more powerful cell lineage tracing methods will help to solve the interesting question of the homology or convergence of respiratory pigment producing cells.

## Conclusion

We propose a new nomenclature for the multigenic family of Metazoan globins based on our phylogenetic study. This phylogeny provides a new scenario for globins and blood evolution: a set of five globin genes, which we call “stem-globins” were present in *Urbilateria*. These genes derived respectively into 5 families of globin genes present in the extant Bilaterians, which we name clade I to V. In the course of Bilaterian speciations, some of these members were lost, others duplicated into paralogous subfamilies. Membrane bound globins are found in all globin clades, except from the clade I. All the known circulatory globins, i.e. globins entering the composition of the respiratory pigments in red blood in Bilaterians, derive from this ancestral clade I stem-globin. We note that the recruitment for the respiratory function of this clade I globin was each time associated with multiple duplication events. The Annelid *Platynereis* has retained the complete set (clade I to V) of ancestral Bilaterian globin genes plus a spiralian-specific globin. *Platynereis* may be an excellent model in the future to determine the initial functions of these stem globins, for instance by selective CRISPR-Cas9 inactivation. It possesses nine clade I globins which are all extracellular. This is consistent with the known presence in various Annelids of the large hexagonal bilayer extracellular hemoglobins with respiratory function. Although the presence of HBL in *Platynereis* has not been formally proven by crystallography, our results confirm it through phylogenetic, expression and morphological evidence. We identified the cells that secrete these extracellular globins. Their location and activity depends on the worm size and life stages. These cells are part of the meso-epithelial walls of some particular vessels: in the trunk in juveniles, and then in the appendages in the pre-mature worms. The blood production level peaks at the onset of the maturation process, likely in preparation for the adult locomotory activity peak. The hemoglobin-producing cell morphology established with EM are similar to pigment producing tissues described in other Annelids but their location in the appendages vessels serving as gills is original.

The finding of hemoglobin-producing cells within the gill organs in a marine animal is an important step forward. *Platynereis* is easily tractable to molecular biology experiments and can be used for obtaining transgenic strains. The molecular characterization of these cells will be very useful for comparative studies and exploring the diversity of hemoglobin-producing cells in Bilaterians. One possibility would be to use single-cell RNAseq, as these hemoglobin-producing cells should be easily singled out in the data because of their massive production of extracellular globins. However, scRNAseq remains relatively expensive and shallow. Another possibility would be to develop a CRISPR-Cas9 protocol for tagging an extracellular globin gene with a GFP coding sequence. This would allow sorting hemoglobin-producing cells with flow cytometry and the bulk sequencing of their transcriptomes.

Another goal will be to understand the embryonic origin not only of the hemoglobin-producing cells but also of the other mesoepithelial cells that are involved in forming the blood vessels. Is there a developmental and lineage connection between these mesoepithelial cells of the vessels, the hemoglobin-producing cells and also the coelomocytes of *Platynereis*? In other words, do we have the equivalent of a hemangioblast lineage in the Annelid? This question is all the more important if we consider the hypothesis that the BVS evolved in the first place to distribute nutrients and was later on recruited for a gas exchange function.

## Methods

### Animal culture and collection

*Platynereis* embryos, juveniles and adults were bred in the Institut Jacques Monod according to the protocols compiled by A. Dorresteijn (www.platynereis.de).

### Survey of *Platynereis dumerilii* globin genes

*Platynereis dumerilii* globin genes were identified with consecutive steps of sequence similarity searches. We first used concatenated human globin sequences as query against expressed sequence tags (ESTs) from *Platynereis* Resources (4dx.embl.de/platy/), and Jékely lab transcriptome (http://jekely-lab.tuebingen.mpg.de/blast/#). Complete coding sequences were assembled from EST fragments using CodonCode Aligner (CodonCode Corporation, USA) and consensus sequences were determined due to high level of polymorphism in this species. The sequences were reciprocally blasted against Genbank non redundant protein database. This allowed constituting a first list of sequences that were recognized as *bona fide* globins. In order to get an exhaustive repertoire of *Platynereis* globin genes, including highly divergent sequences and sequences that might be missing from the transcriptomes, we performed a tblastn search against the most recent *Platynereis* genome assembly using as query a concatenation of all *Platynereis* transcriptome globin sequences. Exons recovered with this method were systematically reblasted on the transcriptome sequences (Additional file 2). No additional gene sequence was detected at this stage. Double or multiple scaffold hits for some gene sequences likely correspond to divergent haplotypes of the same genomic region, the *Platynereis* strain used for genome sequencing showing some highly polymorphic chromosomal regions. Chromosomal globin clusters were annotated using Artemis (http://sanger-pathogens.github.io/Artemis/Artemis/) and for each *Platynereis* globin, intron positions were mapped on genomic DNA using CodonCode Aligner.

### Survey of globin genes in animal genomes

Gene searches were carried out using the tblastn or blastp algorithms implemented in ngKlast (Korilog V 4.0, Questembert, France). We used a concatenation of the *Platynereis* globins as a query. The sequences were reciprocally blasted against Genbank non redundant protein database. This concatenated query sequence covers as much as the molecular diversity of globins as possible and even very derived sequences were recovered in the hit lists. Also, a concatemer query gives a clear drop in the list of hits with decreasing E-values, when sequences with no significant similarity with globins are reached. We took great care however of stopping the screen only when at least ten non-globin proteins turn up in the decreasing E-value list. We used publicly available files of peptide predictions from 22 genome datasets widely distributed among the Metazoan phyla: Porifera, Cnidaria, Placozoa, Ecdysozoa, Lophotrochozoa and Deuterostomia (additional file 13). When available, the screen was complemented with publicly available transcriptomes that allowed for the correction of a few annotation problems. In addition, the globin genes search was performed on the transcriptomes of *Lumbricus terrestris, Arenicola marina* and *Alvinella pompejana*.

The putative extracellular globin genes were identified using Phobius signal peptide predictor (http://phobius.sbc.su.se/, Stockholm Bioinformatics Center) and SignalP-5.0 predictor (http://www.cbs.dtu.dk/services/SignalP/) (additional file 6).

The putative N-terminal acylation predictions were carried out using GPS-Lipid predictor (http://lipid.biocuckoo.org/, Xie et al, 2016) and GPS-Palm predictor (http://gpspalm.biocuckoo.cn/, Ning et al., 2020) and compiled in additional file 6.

The hexagonal bilayer linker proteins were screened in all genomes in the same fashion as globins, using this time as a probe a concatenation of all linker proteins from the earthworm *Glossoscolex* found in Genbank.

### Phylogenetic analyses

The amino-acid sequences of the identified globin genes in 25 species were aligned with MUSCLE 3.7 (66) as implemented on the LIRMM web (http://phylogeny.lirmm.fr/phylo_cgi/one_task.cgi?task_type=muscle) under default parameters and adjusted manually in Bioedit. A selection of aligned positions was produced to eliminate unaligned or ambiguously aligned regions (additional file 4). This resulted in a alignment of 275 globin sequences displaying 231 phylogeny informative positions. The phylogenetic trees were constructed using two different approaches: the maximum likelihood (ML) and the Bayesian analyses. Maximum likelihood trees were generated using PhyML3.0 (http://phylogeny.lirmm.fr/phylo_cgi/one_task.cgi?task_type=phyml) (67), using the LG model of amino-acid substitutions (68). This model has been shown to perform better than the more widely used WAG model. The classical Bootstrap test for short sequences such as globins is inappropriate. Therefore statistical support for nodes was assessed using SH-like test (aLRT score) (69). Bayesian analysis was performed with MrBayes 3.2.6 (70) using either LG or WAG fixed model, run for respectively 32 174 500 generations and 24 972 000 generations, using all compatible consensus and a burn’in value of 0.25. The calculation was performed with 4 chains including one heated chain at temperature 0.5. The average standard deviations of split frequencies were respectively 0.031 and 0.017.

Bayesian posterior probabilities were used for assessing the confidence value of each node. Phylogenetic trees were visualized and rooted using FigTree V.1.5.0 (http://tree.bio.ed.ac.uk/software/figtree/).

### Cloning of extracellular globin genes and in situ probes design

Large cDNA fragments, encompassing at least the whole coding sequences, were cloned by nested PCR using sequence-specific primers on cDNA from mixed larval stages and posterior regenerating segments. PCR products were cloned into the PCR2.1 vector following the manufacturer’s instructions (Invitrogen, France) and sequenced. The full list of primers used is provided in additional file 14. Sequences were deposited in Genbank (MT701024-MT701042). These plasmids were used as template to produce Dig RNA antisense probes for whole-mount *in situ* hybridization (WMISH) using Roche reagents.

### Visualization of *Platynereis* extracellular globin genes expression patterns by whole mount *in situ* hybridization

The animals were fixed in 4% paraformaldehyde (PFA), 1 x PBS, 0.1% Tween20 and stored at −20°C in methanol 100%.

NBT/BCIP whole-mount in situ hybridization was performed as previously described (https://www.ijm.fr/fileadmin/www.ijm.fr/MEDIA/equipes/Balavoine/IJM_HybridationInSituNBT_BCIP.doc) on larval stages (24h, 48h, 72hpf), juvenile larvae, different stage of maturing worms and mature epitoke worms, as well as on regenerated posterior "tails", 9 days after posterior amputation. Bright-field images were taken on a Leica DM5000B microscope equipped with color camera.

### Quantitative PCR

Total RNA was extracted using RNAeasy kit (Qiagen). The extraction was done from a pool of 48hpf larvae, a pool of 6 weeks worms, whole 50 segments juvenile worms, a stretch of 20 segments starting 24 segments after the head for maturing stages. The tissue was disrupted and homogenized inside RLT buffer. First-strand cDNAs were synthesized using 100ng of total RNA, random primers and the superscript II Reverse Transcriptase (Invitrogen, Life Technology). Specific primer pairs were designed with Applied Biosystems Primer Express software to amplify specific fragments between 50 and 60bp (additional file 15). The composition of the PCR mix was: 4uL of diluted cDNA, 5uL of SYBR GREEN master mix, 1 uL of primers fwd, 1uL of primers reverse, in a 10uL final volume. qPCR reactions were run in 96-well plates, in real-time Applied Biosystem StepOne thermocycler. The PCR FAST thermal cycling program begins with polymerase activation and DNA denaturation at 95 °C for 20 sec, followed by 40 cycles with denaturation for 3s at 95°C and annealing/extension for 30s at 60°C. After amplification, melting curve analyses were performed between 95°C and 60°C with increase steps of 0.3 °C to determine amplification product specificity. The slopes of he standard curves was calculated and the amplification efficiencies (E) were estimated as E=10Λ(−1/slope). qPCR were run with biological triplicates for juvenile worms of 50 segments and maturing worms, and each sample was run with technical duplicates and the ribosomal protein small subunit 9 (rps9) and *D-adenosylmethionine synthetase* (sams) housekeeping gene. Relative expression level of each target gene was obtained by 2Λ(−ΔCt sample) where ΔCt sample = Ct sample-Ct gene of reference (average of rps9 and sams).

### Transmission electron microscopy

For electron microscopy animals were relaxed in 7,5% MgCl2, fixed in 2,5% glutaraldehyde buffered in 0,1M phosphate buffer and 0,3M NaCl, rinsed 3 times in the same buffer, and postfixed in 1%OsO4 in the same buffer for 1h. The specimens were dehydrated in ascending acetone series, transferred in propylene oxide and embedded in Epon-Araldite resin (EMS). Ultrathin sections (60-80nm) were cut with Reichert Ultracut E or Leica Ultracut UCT and counterstained with 2% uranyl acetate and Reynolds lead citrate. Images were acquired using Zeiss Libra 120, FEI Technai or Jeol JEM-1400 transmission Electron microscopes and processed with Fiji and Adobe Photoshop software. For 3D-reconstructions the series of semithin (700 nm) sections were cut with Diatome HistoJumbo diamond knife (Blumer et al., 2002), stained with methylene blue/basic fuchsine (D’amico, 2009), and digitalized at 40x magnification using Leica DM2500 microscope with camera. The images were aligned with IMOD and ImodAlign tool. The reconstructions were made with Fiji TrackEM2 plugin.

## List of abbreviations

BVS: blood vascular system
RBC: red blood cells
qPCR: quantitative polymerase chain reaction

## Additional information

**Additional file 1: Amino acids alignment of the** *Platynereis* **extracellular globins**with the respiratory globins of *Arenicola marina*, *Lumbricus terrestris* and human hemoglobins. The highly conserved residues of globins in dark red (75). The conserved cysteines NA2 and H7 residues known to be involved in the formation of an intrachain disulfide chain in Annelids are shown in black (28).

**Additional file 2: Table of the** *Platynereis* **globin blasted on the** *Platynereis dumerilii* **genome assembly.**Sheet 1: the hits for each mRNA sequence are listed in order of decreasing E-Value. Each hit partial sequence was reciprocally blasted on the Platynereis mRNAs and on Genbank for unidentified sequences. Some of the hits correspong to likely distant haplotypes. Others do not correspond to a globin sequence. Sheet 2: the genome assembly was screened with a concatenated *Platynereis* transcriptome globin probe to detect potential additional genes. The hits are listed by decreasing E-value. Partial hits were reciprocally blasted on Genbank and Platynereis transcriptome. Most high E-value hits correspond to the genes already identified in the trabscriptome. A series of low E-value, short sequence globin hits on Genbank were found to code for *Platynereis* proteins that bear no similarity to globins, indicating that no additional globin gene is present in the assembly.

**Additional file 3: List of all globin predicted protein sequences used in this study; list of hexagonal bilayer linker predicted protein sequences identified in all genomes.**

**Additional file 4: Nexus file of aligned globin sequences used for the phylogenetic analysis**. The unaligned sequences at the N- and C-termini were cropped as well as some insertions which likely correspond to undetected intronic sequences.

**Additional file 5: Maximum likelihood tree using LG model and a SH-like test of a dataset of 272 sequences from 24 metazoan species, (simplified in Figure 3).**The letter and color codes for species names is explained at the right of the figure. The tree is arbitrarily rooted outside of the clear natural groups. The tree shows 5 groups (clade I, clade II, clade III, clade IV, clade V), 4 of the 5 with well-supported nodes (clade I, clade II, clade III, clade IV), encompassing sequences of most Bilaterian phyla. An additional well-supported node groups together only Lophotrochozoan species sequences. Support values for maximum-likelihood with LG model, bayesian posterior probability with LG model and bayesian posterior probability with WAG model are indicated on the key nodes discussed. *Platynereis* sequences are indicated in red labels. The globin peptides predicted to be extracellular or membrane-bound are indicated with blue or green rectangles, respectively.

**Additional file 6: Table of all globin sequences indicating their clade affinity in the ML tree, the signal peptide prediction and N-terminal acylation prediction.**Signal pepetide prediction is done with SignalP 5.0. Myristoylation is predicted with GPS-lipid algorithm and palmitoylation with GPS-Palm algorithm. The dual acylation prediction is based on two closely located myristoylation and palmytoylation sites only.

**Additional file 7: List of additional globin protein sequences,**from non Bilaterian Metazoans (*Aurelia aurita*, *Clytia hemisphaerica*, *Hydra vulgaris*, *Amphimedon queenslandica*, *Mnemiopsis leydyi*), one Acoel (*Symsaggitifera roscoffensis*), and two Choanoflagellates (*Salpingoeca rosetta*, *Monosiga brevicollis*). The two globins from the Acoel are extracted from a transcriptome (7). Acoels are part of Xenacoelomorpha, a group of worms that has been proposed to be the sister-group of all remaining Bilaterians (76). This view remains however challenged as they are also proposed to belong with the Deuterostomes (77). Choanoflagellates are a group of Eukaryots closely related to Metazoans, that had been reported to possess neuroglobin-like proteins (7).

**Additional file 8: Maximum likelihood tree using LG model and a SH-like test of a dataset of 293 sequences from 32 metazoan species.**The tree is arbitrarily rooted outside of the natural clades. Green and red diamonds indicate a number of nodes that are supported by aLRT values superior to 0.75 and 0.95, respectively. The six clades of globins previously highlighted are again recovered. as Hydrozoans (in our analysis, *Hydra* and *Clytia*) possess genes related to the clade III. One Ctenophore, *Mnemiopsis*, also display a clade III related gene. One occurrence of a clade I gene in the Hydrozoan *Clytia* appears suspicious as the protein does not show any particular similarity with clade I globins in a blast search. Last, the Acoel is shown to possess clade III and V globins. In the absence of a complete genome sequence and the phylogenetic position of Acoels being ambiguous, it does not inform on the early evolution of globins.

**Additional file 9: The life cycle of** *Platynereis dumerilii*. *Platynereis dumerilii* is a medium-sized Annelid that can be easily cultured in the laboratory, giving a large number of offspring all year round. The life cycle is fairly typical of marine Annelids. It includes a microscopic (160 μm diameter) lecithotrophic trochophore larva that elongates after 2,5 days in a minute three-segment worm (400 μm long). Once settled in the benthos in a silk tube, the larva will grow by posterior addition of segments and in cross-section to reach a considerably larger size (5-6 cm). The worm lives a relatively sedentary life, feeding on a variety of fresh or decaying food in the benthos for the longest part of its lifespan. Nearing the end of its life, the worm undergoes a rather dramatic sexual metamorphosis: the coelom, from head to tail, entirely fills up with gametes; the worm segmental appendages, called parapodia, change shape and acquire a swimming locomotory function; the gut degenerates as adults do not eat. Both males and females exit their tubes to swarm at the surface of the sea. At this stage they acquire a very fast swimming behaviour and have only a few hours for mating before they die of exhaustion.

**Additional file 10: Expression patterns of five extracellular globin genes in** *Platynereis dumerilii.* WMISH on juvenile stages (30-35 segments). The expression patterns of all extracellular globins are located in the same hemoglobin-producing cells of transverse trunk vessels (red arrowheads) and parapodial vessels (blue arrowheads).

**Additional file 11: Movie of a 3D reconstruction of the vasculature of a 50-segment sub-adult worm.** 3 consecutive segments are shown.

**Additional file 12: Table of relative normalized expression of *Platynereis* extracellular globins measured by qPCR.**

**Additional file 13: Table of genomic and transcriptomic resources used for 32 species.**

**Additional file 14: List of primers used for PCR amplification of *Platynereis* globin cDNAs.**

**Additional file 15: List of primers used for qPCR amplification of *Platynereis* extracellular globins.**

## Declarations

### Ethics approval and consent to participate

“Not applicable” --- This research focuses on non cephalopod invertebrates. There are no ethical considerations mentioned for these species according to EU Directive 86/609-STE123.

### Consent for publication

"Not applicable"

### Availability of data and materials

The data sets supporting the results of this article are available in the Figshare repository (https://figshare.com/account/home#/projects/72548).

### Author's contributions

SS performed all molecular biology experiments, analysed the data, composed the figures. GB conceived the study, analysed the data, composed the figures and drafted the text. VVS performed the semi-thin and TEM imagery, analysed the data and composed the figures. XB participated in the phylogenetic analyses. PK participated in the genomic and transcriptomic screenings. AJMC and CR participated in data analysis. All authors corrected the draft. All authors read and approved the final manuscript.

## Acknowledgements

We warmly thank Detlev Arendt's laboratory (EMBL Heidelberg, Germany) and Oleg Simakov's laboratory (University of Vienna, Austria) for letting us use the data of the *Platynereis* whole genome sequencing and assembly ahead of publication. We thank the Balavoine Lab Members and Biofluidics Lab members for helpful discussions during the course of this work. We acknowledge Nathalie Luciani for equipment and training for qPCR experiments and analysis. We acknowledge the ImagoSeine facility (member of the France BioImaging infrastructure supported by the French National Research Agency, ANR-10-INSB-04, ‘Investments of the future’) and its experienced staff for their assistance in imaging and image analysis.

