## Supplementary figures and images for "Globins in the marine Annelid *Platynereis dumerilii* shed new light on hemoglobin evolution in Bilaterians"

### Additional file 1

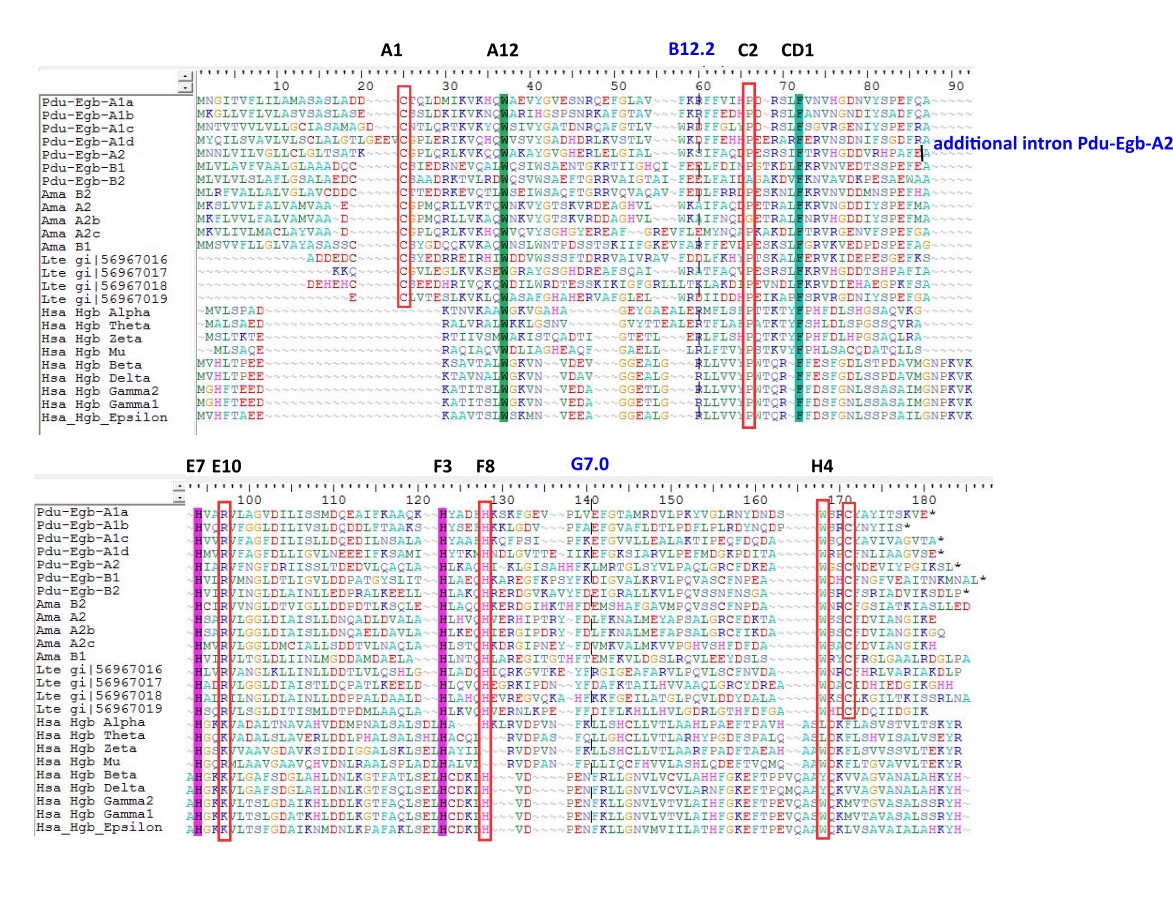

### Additional file 5

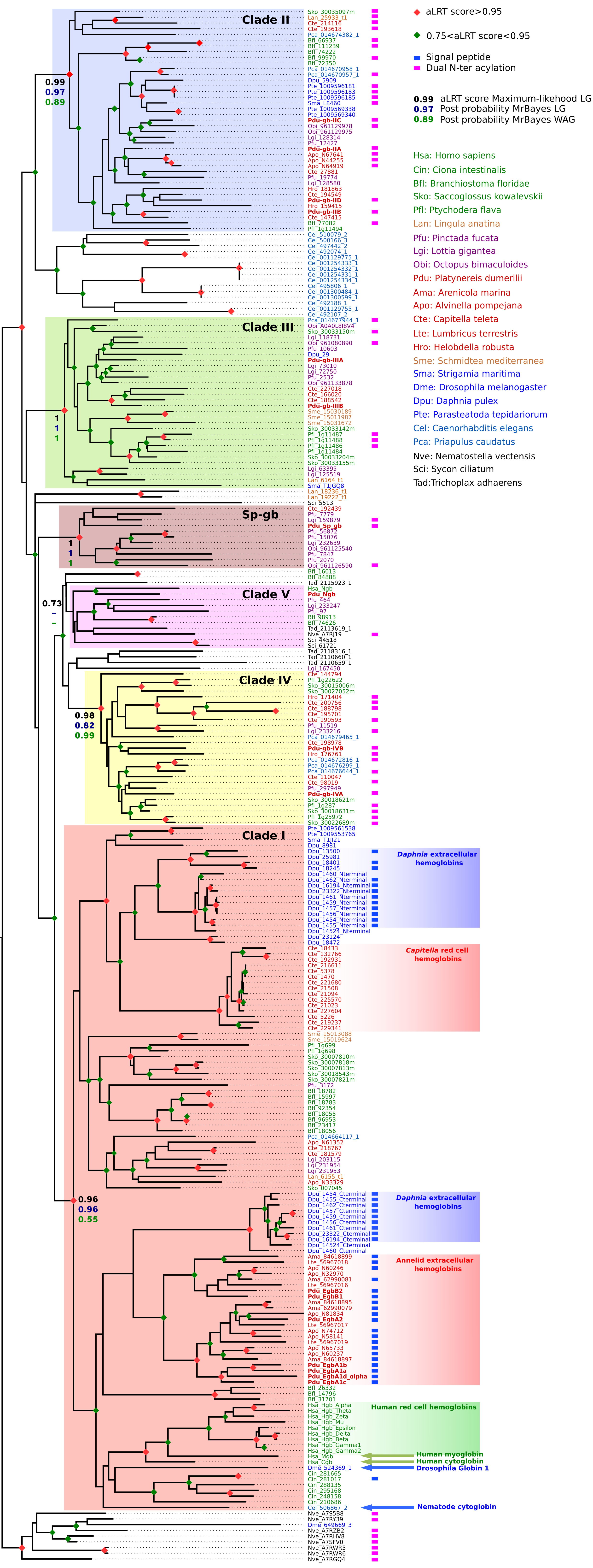

### Additional file 8

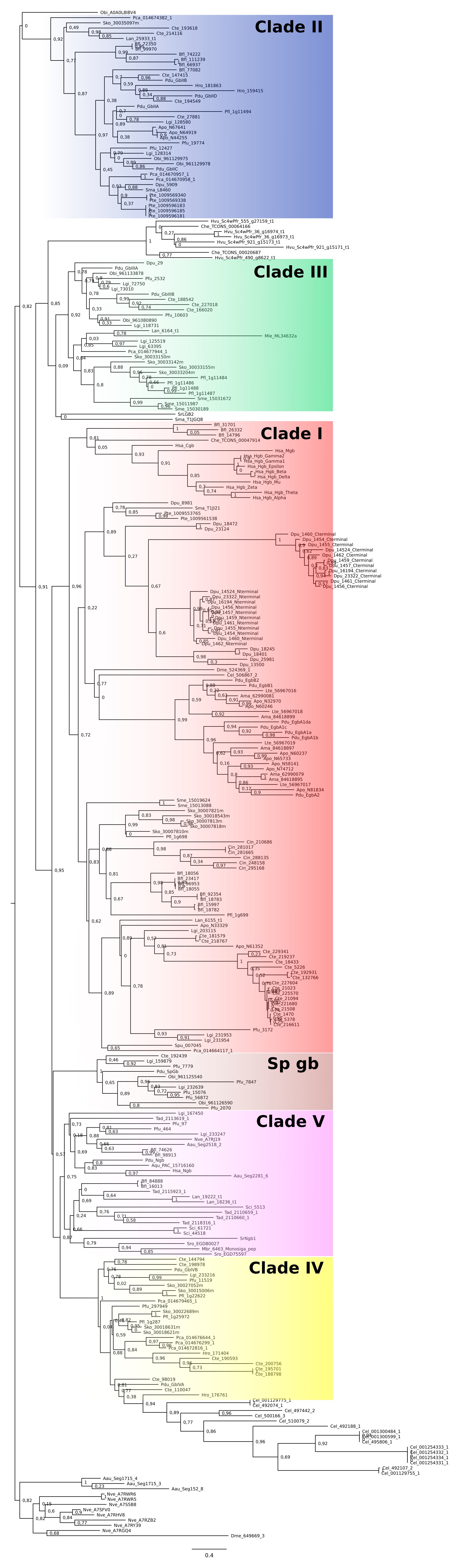

### Additional file 9

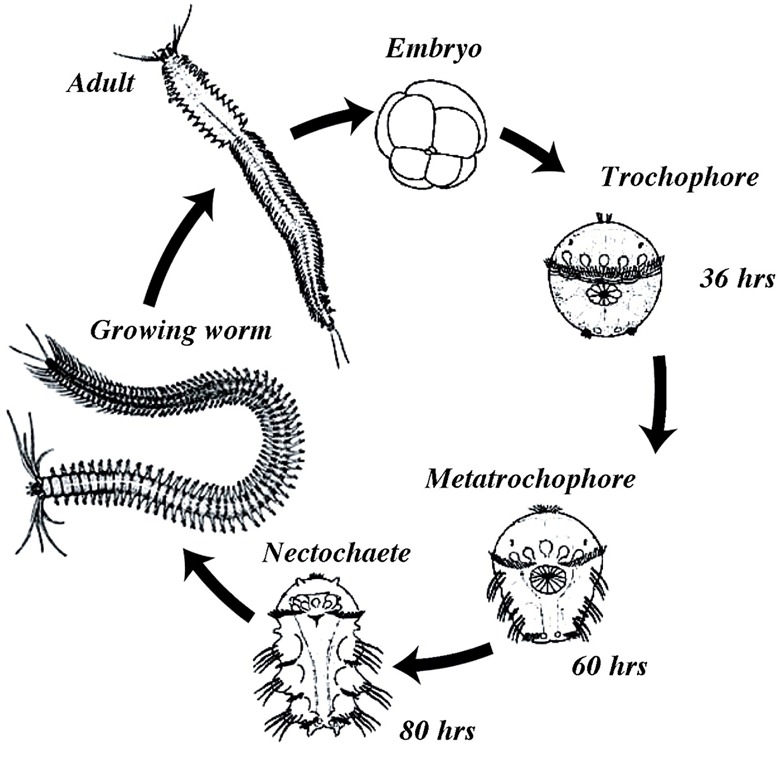

### Additional file 10

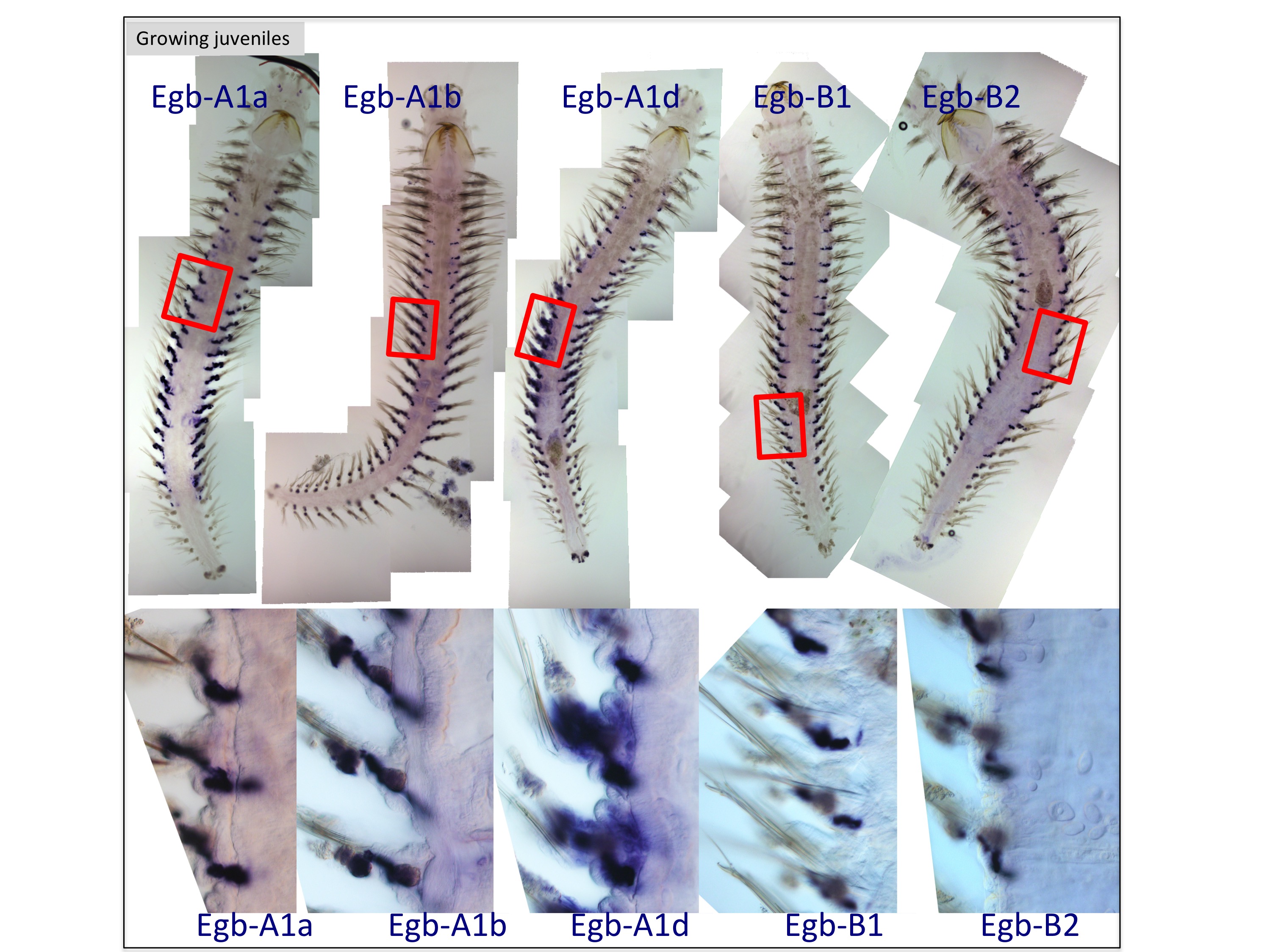
