## Additional file 14 for "Globins in the marine Annelid *Platynereis dumerilii* shed new light on hemoglobin evolution in Bilaterians"

>Pdu_Egb_A1a_ F1
CGCCCCAATACTGGTAACTG

>Pdu_Egb_A1a_ R1
TTTTGCAACCAAGCCACATA

>Pdu_Egb_A1a_ F2
TGGCTGATTCAAGATGAACG

>Pdu_Egb_A1b_F1
GAGCCAATCACCTTCTGGTC

>Pdu_Egb_A1b_R1
TTTACAGTGCTCGCTCATGG

>Pdu_Egb_A1b_F2
CCAATCACCTTCTGGTCCTG

>Pdu_Egb_A1c_F1
CAGAGTACGCGGGGCTTC

>Pdu_Egb_A1c_R1
GTCGCCACACCAAAACCAAA

>Pdu_Egb_A1c_F2
GGCTTCAGACCATCTCGAGG

>Pdu_EgbA1c_R2
CGCCACACCAAAACCAAAGAT

>Pdu_Egb_A1d-alpha_F1
GATCAAGGTCCAGCACCAGT

>Pdu_Egb_A1d-alpha_R1
TGCTTATGAACAATGAGGACCA

>Pdu_Egb_A1d-alpha_R2
TGAACAATGAGGACCATACAGG

>Pdu_Egb_A2 _F1
CTGACTTCTGCCACCAAGTG

>Pdu_Egb_A2_R1
GGTCTGGTGAAAAGCCAAGA

>Pdu_Egb_A2_F2
CTGAAGGTCAAGCAGCAGTG

>Pdu_Egb_B1_F1
ATTTGCCACTAGGGGAAGGA

>Pdu_Egb_B1_R1
CACAAAGCGATCAACACAGG

>Pdu_Egb_B1_F2
TAGGGGAAGGAAGGGATGTT

>Pdu_Egb_B2_F1
GAGTACGCGGGGAGTCAATA

>Pdu_Egb_B2_R1
 CTTCGCCGGACTGAAGAGTA

>Pdu_Egb_B2_F2
CCCTAGCTTTCTTGGGCTCT

>Pdu_Egb_B2_R2
 CTGGGGAACGTTTGAACAAT

>Pdu_Ngb_F1
TTTGCGATTCGGAGTTGGGA

>Pdu_Ngb_R1
AGACAATATCCCTAACCCACACA

>Pdu_Ngb_F2
CGATTCGGAGTTGGGAGTCA

>Pdu_Ngb_R2

AGGTACATTCCCTTAAAATCCCA

>Pdu_gb_IIIB_F1

ACATCGTCCAAAGGTCAAGG

>Pdu_gb_IIIB_R1

TGACACTCCAGAAGGTGACG

>Pdu_gb_IIIB_F2

CGTCCAAAGGTCAAGGAAAA

>Pdu_gb_IID_F1

TAAAGATGTGGCCCTGGAAC

>Pdu_gb_IID_R1

AGGCCAATGTATGGCTTCAC

>Pdu_gb_IID_F2

CATCCGGTTCATCCTCAACT

>Pdu_gb_IVA_F1

TAAAGCGGACGCGAAAGTAT

>Pdu_gb_IVA_R1

TGGAAGGTCAATGTGGGAGT

>Pdu_gb_IVA_R2

TGCGCAGAAGTGTTTTTGTC

>Pdu_gb_IVB_F1

GGCCAGGATAGCAACCATAA

>Pdu_gb_IVB_R1

GCTGCCCAGAAGCAAAATAA

>Pdu_gb_IVB_F2

CCATAAAAATGCCGACCAAC
