## Additional file 15 for "Globins in the marine Annelid *Platynereis dumerilii* shed new light on hemoglobin evolution in Bilaterians"

qPCR primers for Platynereis dumerilii extracellular genes

>qPCR-Pdu-EgbA1a-F

TGAGAGACGTTCTGCCCAAA

>qPCR-Pdu-EgbA1a-R

GGCTCCAGGAATCGTTATCG

>qPCR-Pdu-EgbA1b-F

AAAGTTGGGAGATGTGCCATTT

>qPCR-Pdu-EgbA1b-R

CGGGCAAAGTGTCCAAGAA

>qPCR-Pdu-EgbA1c-F

CTCTGCTCTTGCCCATTACG

>qPCR-Pdu-EgbA1c-R

TTCCTTGAAGGGAATGCTTG

>qPCR-Pdu-EgbA1d-F

GAAGAGCGTGCACGTTTTGAG

>qPCR-Pdu-EgbA1d-R

GTCTCCAGAAAAAATGTTGTCG

>qPCR-Pdu-EgbA2-F

CGGCTTCGACAGAATCATCTC

>qPCR-Pdu-EgbA2-R

CCAGTTGGGCCTGAAGGA

>qPCR-Pdu-EgbB1-F

CAGGCTCTTTGGCAATCCA

>qPCR-Pdu-EgbB1-R

GCCGATGATGGTTCTCTTTCC

>qPCR-Pdu-EgbB2-F

AGCTGCCGATAGGAAGACC

>qPCR-Pdu-EgbB2-R

GTGAACTCAGCGGACCAGA

>qPCR-Pdu-cdc5-F

CCTATTGACATGGACGAAGATG

>qPCR-Pdu-cdc5-R

TTCCCTGTGTGTTCGCAAG

>qPCR-Pdu-rps9-F

CGCCAGAGAGTTGCTGACT

>qPCR-Pdu-rps9-R

ACTCCAATACGGACCAGACG

>qPCR-Pdu-sams-F

CAGCAACGGTGAAATAACCA

>qPCR-Pdu-sams-R

CATCACTCACTTGATCGCAAA
